# Novel insights in the pathophysiology of α-synuclein dysregulation on D2 receptor activity contributing to the vulnerability of dopamine neurons

**DOI:** 10.1101/2021.03.30.437775

**Authors:** Abeer Dagra, Douglas R. Miller, Fatemeh Shaerzadeh, Min Lin, Adithya Gopinath, Sharonda Harris, Zachary A. Sorrentino, Sophia Velasco, Adetola R Alonge, Janelle Azar, Joe J Lebowitz, Brittany Ulm, Anthea-Mengfei Bu, Carissa A. Hansen, Nikhil Urs, Benoit I. Giasson, Habibeh Khoshbouei

## Abstract

Pathophysiological damages and loss of function of dopamine neurons precedes their demise and contributes to the early phases of Parkinson’s disease. The presence of aberrant intercellular pathological inclusions of the protein α-synuclein within ventral midbrain dopaminergic neurons is one of the cardinal features of Parkinson’s disease. We employed multiple complementary approaches in molecular biology, electrophysiology, and live-cell imaging to investigate how excessive α-synuclein levels alters multiple characteristics of dopaminergic neuronal dynamics and dopamine transmission prior to neuronal demise. These studies demonstrate that α-synuclein dysregulation of D2 receptor autoinhibition contributes to the vulnerability of dopaminergic neurons, and that modulation thereof can ameliorate the resulting pathophysiology. These novel findings provide mechanistic insights in the insidious loss of dopaminergic function and neurons that characterize Parkinson’s disease progression with significant therapeutic implications.

## Introduction

Volitional movement is a crucial fundamental behavior of everyday life that is often taken for granted until control diminishes and deteriorates. Dopaminergic neurons within the ventral midbrain play a critical role in the initiation and control of volitional movement^1,2^ and the progressive demise of these neurons is a defining hallmark of Parkinson’s disease (PD)^3^. Pathophysiological damages and loss of function of these neurons precedes their demise and contributes to the early phases of the movement impairments^4^. No current therapies reverse or ameliorate the insidious nature of PD or the many related neurodegenerative diseases associated with the demise of dopaminergic neurons. The lack of a better understanding of the etiology of PD greatly impedes the ability to identify therapeutic targets to slow this detrimental progression.

The presence of aberrant intercellular, pathological inclusions comprised of the protein α-synuclein (α-syn) in the form of Lewy bodies and Lewy neurites within ventral midbrain dopaminergic neurons is another cardinal feature of PD^5–7^. Several missense mutations in the α-syn gene *(SNCA)* can be responsible for familial PD, but the duplication^8^ or triplication^9^ of *SNCA* are also sufficient to cause PD and the related disease Lewy body dementia. Thus, only a 50% increase in the expression of wild type α-syn as in the duplication of the *SNCA* gene is sufficient for detrimental outcome on dopaminergic neurons resulting in disease. Furthermore, some studies indicate that elevated α-syn levels also occurs in idiopathic PD, but the pathophysiological mechanisms associated with increased levels of α-syn remain poorly understood.

In the studies, herein, extensive complementary approaches in molecular biology, electrophysiology, and live-cell imaging were utilized to investigate the effects and outcomes of α-syn elevated expression on dopaminergic neuronal dynamics, intraneuronal calcium homeostasis and dopamine transmission prior to neuronal demise. It is demonstrated that D2 receptor autoinhibition, contributes to alterations in neuronal homeostatic properties, and that modulation thereof can ameliorate the pathophysiology resulting from excessive α-syn levels. These results provide novel mechanistic insights in the pathobiological impact of α-syn on dopaminergic neuron function and their demise characteristic of the insidious nature of PD.

## Results and discussion

### Tyrosine hydroxylase promoter-driven adeno-associated virus (AAV) efficiently transduces and express human α-synuclein in cultures midbrain dopamine neurons

In order to investigate the pathophysiology changes associated with α-syn overexpression we first develop a cell model with high fidelity expression of human α-syn in midbrain dopaminergic neurons. We utilized a tyrosine hydroxylase (TH) promoter-driven adeno-associated virus (AAV) to specifically express wild-type human α-syn in these cultured dopaminergic neurons. First to demonstrate the specificity of this vector, cultures were transduced with the virus expressing GFP. As demonstrated in Figure 1A-B, 91% ± 3 of neurons expressing GFP-are TH-positive, indicating high specificity. The same pAAV1-TH backbone but with the human α-syn cDNA, was utilized to overexpress human α-syn in dopamine neurons. The transduction of pAAV1-TH-human-αsyn in cultured midbrain dopamine neurons was confirmed via immunocytochemistry and western blot analyses, demonstrating elevated expression of α-syn these neurons (Figure 1C-D, Supplemental Figure 1, p = 0.005, two-tailed t-test, n = 3 independent experiments).

**Figure 1.**
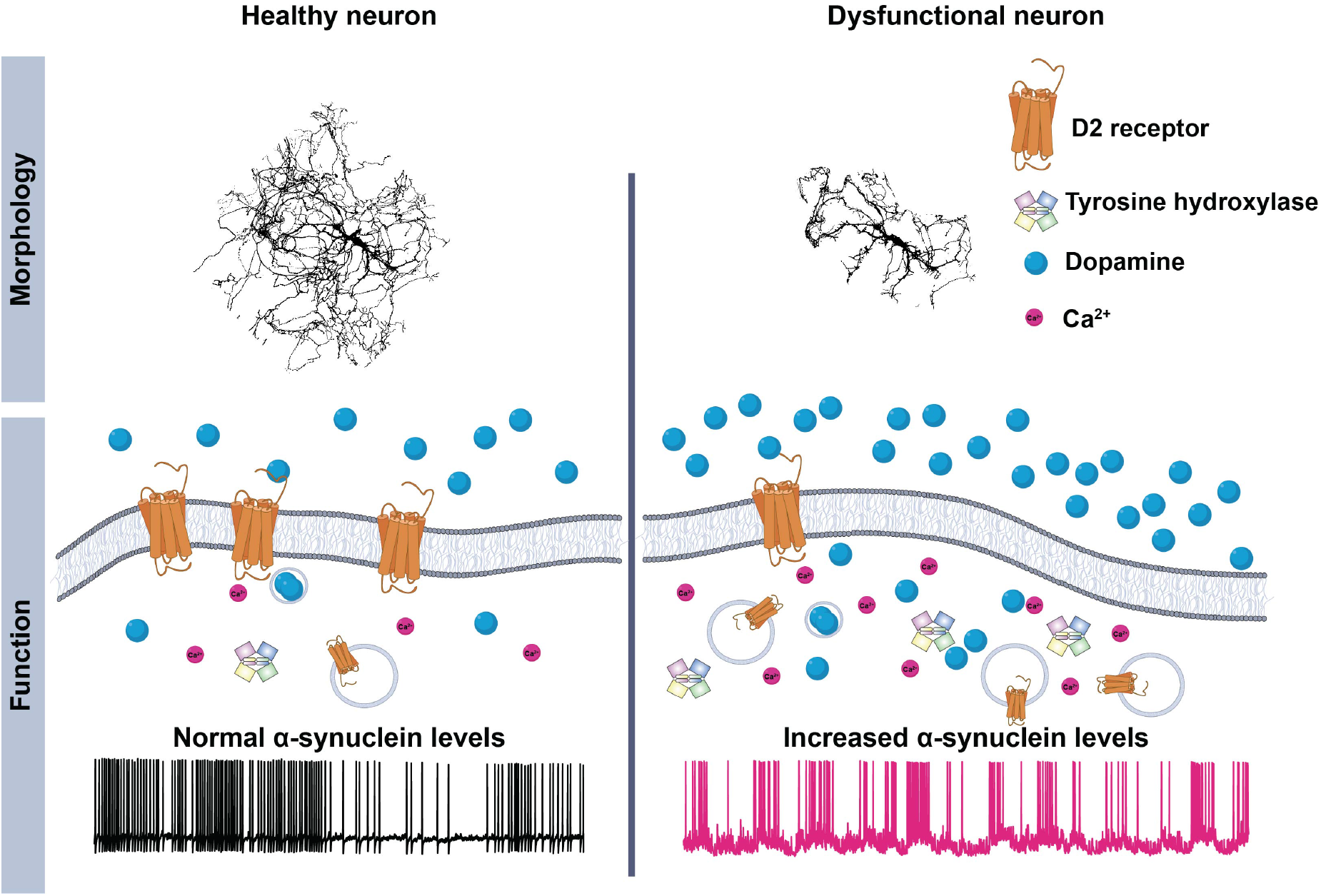
Tyrosine hydroxylase (TH) promoter driven adeno-associated virus (AAV) efficiently transduces human α-synuclein or the control construct (TH-GFP) in midbrain dopamine neurons. (A, B) Immunolabeling of TH confirmed 91% ± 3 of TH positive neurons co-express GFP, suggesting a high fidelity for pAAV1-TH-GFP viral transduction in the TH positive neurons (n = 3 independent experiments). (C, D) The transduction of pAAV1-TH-human-αsyn in midbrain dopaminergic neurons was confirmed via immunocytochemistry analysis and western blot (n = 4 independent experiments, see supplemental figure 1 for Western blot analysis). Scale bars: 50 μm.

### Overexpression of α-synuclein disrupts calcium dynamics and firing activity of dopamine neurons

.Increased α-syn burden in dopamine neurons is correlated with neuronal loss in neurodegenerative diseases such as Parkinson’s disease (PD)^10,11^. Although extensively studied in cortical neurons, yeast, and heterologous expression systems^12–17^, α-syn regulation of intracellular calcium and firing activity in dopaminergic neurons prior to cell death remains less clear. Maintenance of calcium homeostasis is a vital process in neurons^18–20^. Calcium is a ubiquitous second messenger that helps to transmit depolarization status and synaptic activity to the biochemical machinery of a neuron^21^. High intensity calcium signaling requires high ATP consumption to restore basal (low) intracellular calcium levels. Increased intracellular calcium may also lead to increased generation of mitochondrial reactive oxygen species^22,23^. Impaired abilities of neurons to maintain cellular energy levels and to suppress oxygen species may impact calcium signaling during aging and in neurodegenerative disease processes^5,6^. To investigate if α-syn overexpression regulates dopaminergic neuronal activity prior to neuronal demise, we employed live cell calcium imaging recording in DAT-Cre-GCaMP6f-expressing dopamine neurons containing either endogenous levels of α-syn (naive) or overexpressing α-syn. Compared to the modest calcium events in naive dopaminergic neurons, both width and amplitude were increased in the presence of α-syn overexpression, creating repeated burdens on the neuron (Figure 2A-C, width - p = 0.034, one-way ANOVA, F(2,97) = 3.89, n = 33 wild type neurons, n = 40 α-syn overexpressing neurons, amplitude - p = 0.000002, one-way ANOVA, F(2,1294)=12.62, n = 33 wild type neurons, n = 40 α-syn overexpressing neurons). These data suggest that increased levels of α-syn in dopamine neurons leads to disturbances in calcium homeostasis that can alter biophysical properties of neurons, neuronal activity, neurotransmission^20,24–29^ and neuronal death, all of which are shared hallmarks in neurodegenerative diseases^24,30–32^.

**Figure 2.**
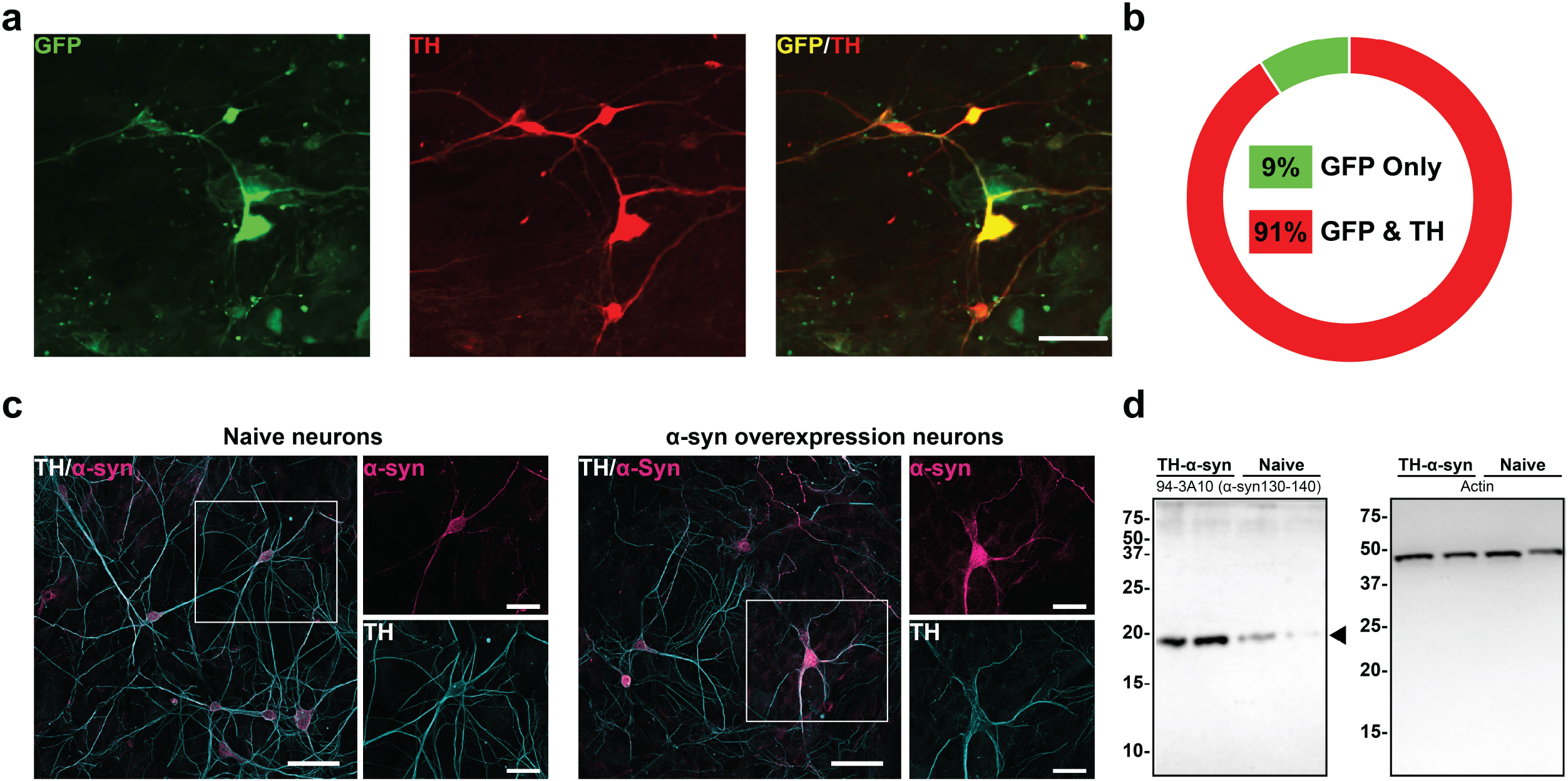
Overexpression of α-synuclein disrupts calcium dynamics and firing activity of dopaminergic neurons. (A) Top - Representative spontaneous GCaMP6f calcium activity in naive dopaminergic neurons (left, black) and dopaminergic neurons overexpressing α-syn (right, pink) exemplify the alteration in calcium dynamics due to increased levels of α-syn. Bottom - Spontaneous calcium activity encompassing all neurons recorded in each experimental group (n = 33 wild type neurons, n = 40 α-syn overexpressing neurons, form 8 biological replicates). (B) Calcium events in all neurons were defined as fluctuations of the fluorescent signal at least two standard deviations above the fluorescence baseline value. (C) Spontaneous calcium events were quantified for event rate, width, and amplitude. Whereas overexpression of α-syn does not alter calcium event rate (p = 0.2596, one-way ANOVA, F(2,65) = 1.38 n = 33 wild type neurons, n = 40 α-syn overexpressing neurons), α-syn burden results in broadening of calcium events (p = 0.034, one-way ANOVA, F(2,97) = 3.89, n = 33 wild type neurons, n = 40 α-syn overexpressing neurons) and increases in amplitude (p = 0.000002, one-way ANOVA, F(2,1294) = 12.62, n = 33 wild type neurons, n = 40 α-syn overexpressing neurons). (D) Representative whole cell current-clamp recordings of spontaneously active naive (left, black) dopaminergic neuron compared with a dopaminergic neuron overexpressing α-syn (right, pink). (E) Naïve dopaminergic neurons fire in a canonical pacemaker pattern and consistent rate, whereas with α-syn overexpressing they fire significantly more frequently (firing frequency: from eight independent experiments, 1.132 ± SEM for naive neurons vs. 3.187 ± SEM for α-syn overexpressing neurons, two-tailed unpaired t-test, p = 0.0142; interspike interval (ISI): 1143 ± SEM for naive neurons compared to 497.9 ± SEM for α-syn overexpressing neurons, two-tailed unpaired t-test, p = 0.0431) and trend in bursts with intermediated periods of quiescence (CV of ISI - 1.676 ± SEM for naive neurons vs. 2.932 for α-syn overexpressing neurons, two-tailed unpaired t-test, p = 0.0808 box plot whiskers represent the 95% confidence interval, the upper and lower bounds of the box represent the 75^th^ and 25^th^ percentiles, respectively, with the middle line indicative of the median value of the sample). Empty circles in panel C represent statistical outliers all of which were included in the analyses. Bar graphs ± SEM are overlaid with individual filled data points.

While measurement of calcium activity provides inferential information about neuronal firing, we did not observe a change in calcium event rates (Figure 2C, left). To directly test whether α-syn overexpression modulates firing activity of dopaminergic neurons, we utilized whole-cell current clamp recordings to measure the spontaneous firing activity of these neurons. While dopaminergic neurons containing an endogenous level of α-syn exhibited the characteristic pacemaker-like firing activity^33,34^, the spontaneous firing activity of α-syn overexpressing dopamine neurons showed an irregular and clustered firing pattern with increased burst firing activity within the clusters (Figure 2 D-E, from eight independent experiments, 1.132 ± SEM for naive neurons vs. 3.187 ± SEM for α-syn overexpressing neurons, one-way ANOVA, followed by Tukey’s HSD, F(2,20) = 6.092, WT vs α-syn p = 0.0123). Thus far, our data suggest that increased α-syn levels in dopaminergic neurons lead to altered calcium dynamics and increased firing activity. Both firing activity and calcium dynamics in dopamine neurons are tightly regulated by the activity of the D2 autoinhibitory receptors^33–37^. Therefore, we asked whether α-syn-induced dysregulation of dopamine neuronal activity and calcium dynamic are due to reduced D2-receptor autoinhibition.

### α-synuclein overexpression reduces D2-receptor autoinhibition

Multiple channels and transporters regulate neuronal activity and intracellular calcium dynamics. Dopamine activation of D2 autoinhibitory receptors on dopamine neurons decreases neuronal activity^33–36,38,39^ and intracellular calcium dynamics^31,35,40–43^. Therefore, we tested the hypothesis that the observed disturbance in neuronal excitability and calcium dynamics in α-syn overexpressing neurons is due to dysregulation of canonical D2 receptor (D2R) autoinhibition in these neurons. In a double-blinded experimental design, we exposed DAT-Cre-GCaMP6f-expressing dopaminergic neurons to dopamine (1 μM) while monitoring the change in GCaMP6f signal. Consistent with the literature^44,45^, naive neurons exhibited the characteristic decreased fluorescence during exposure to extracellular dopamine intensity (Figure 3A,B - top, n = 14-21, two-way ANOVA where the variables are time and treatment, followed by Tukey’s HSD, p = 0.0003, from five independent replicates). However, α-syn overexpressing neurons exhibited attenuation to dopamine-induced reduction of GCaMP6f signal, (n = 17-26, two-way ANOVA where the variables are time and treatment, followed by Tukey’s HSD, p = 0.0088, from five independent replicates). Further, fold change responses of naive dopamine neurons were significantly stronger than α-syn overexpressing neurons (Figure 3C, n = 11-17, one-way ANOVA followed by Tukey’s HSD, WT vs. α-syn, p = 0.0035, WT vs. GFP p = 0.2927 and α-syn vs. GFP p = 0.0001, from five independent replicates, Supplemental Figure 2A). These data suggest α-syn overexpression decreases inhibitory feedback regulation of dopamine neurons of intraneuronal calcium activity.

**Figure 3.**
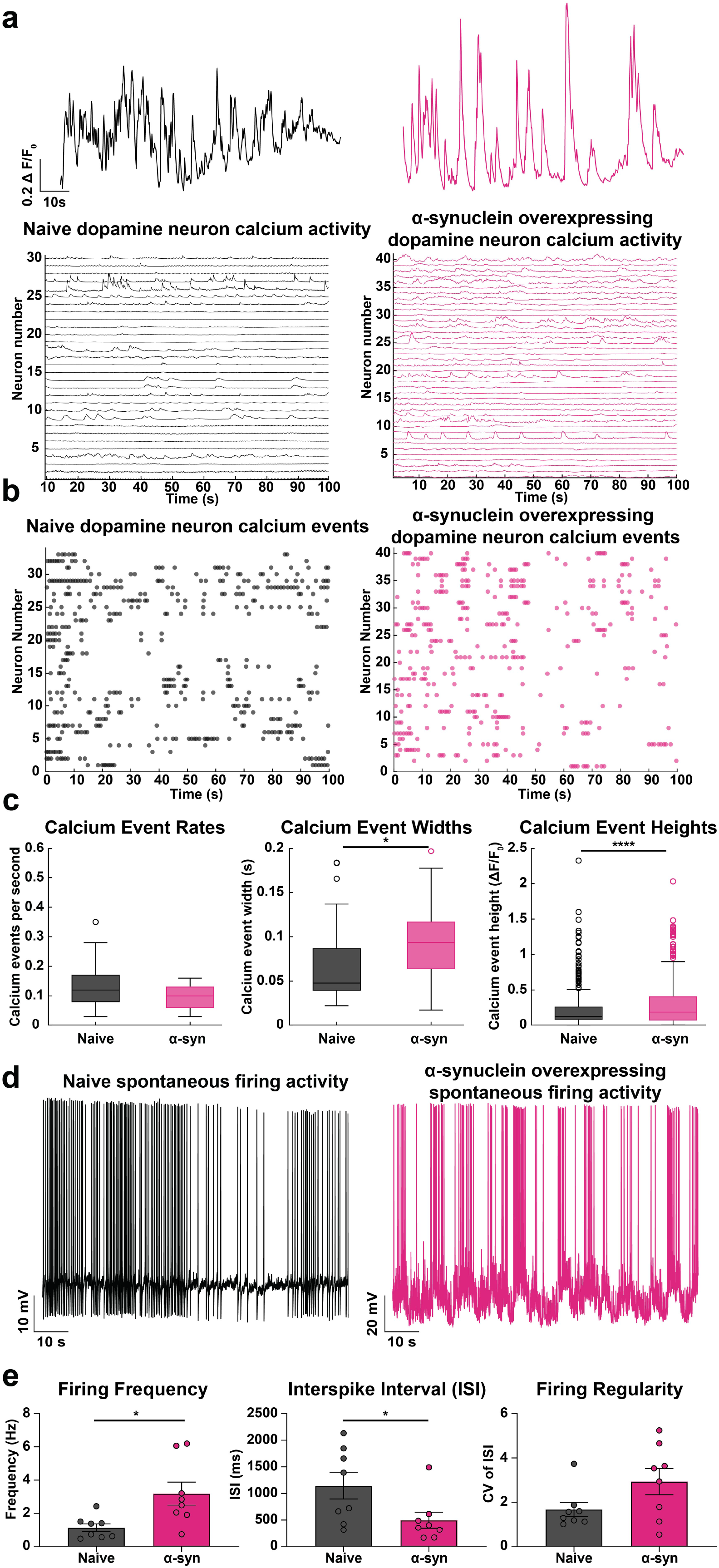
α-synuclein overexpression reduces D2 receptor autoinhibition. Live-cell GCaMP6f calcium imaging in DAT-cre-GCaMP6f-expressing dopamine neurons was employed to measure the peak amplitude of calcium events at baseline (sampling frequency of 1 image/second), and after bath application of dopamine (1 μM) or quinpirole (10 μM) for 150 seconds (sampling frequency of 1 image/second). Peak fluorescence intensities were normalized to average baseline GCaMP6f fluorescence intensity (initial 60 seconds). The dotted line denotes normalized average baseline GCaMP6f fluorescence intensity (30 seconds prior to treatment shown). (A) Top panels - representative images of naive dopaminergic neurons and bottom panels - α-syn overexpressing dopaminergic neurons before (left panel) and during dopamine (1 μM) administration (right panels). (B) Top - In naive dopaminergic neurons, bath application of dopamine rapidly and significantly reduced the GCaMP6f fluorescence intensity (n = 14-21, two-way ANOVA where the variables are time and treatment followed by Tukey’s HSD, p = 0.0003, from five independent replicates). Bottom - In α-syn overexpressing dopaminergic neurons, bath application of dopamine produced a smaller reduction in the peak amplitude of calcium events as indicated by decreased GCaMP6f fluorescence intensity (n = 17-26, two-way ANOVA where the variables are time and treatment followed by Tukey’s HSD, p = 0.0088, from five independent replicates). (C) The fold change in the average GCaMP6f fluorescence intensity before and after drug administration. Dopamine-mediated autoinhibition response is reduced in α-syn overexpressing neurons (n = 11-17, one-way ANOVA followed by Tukey’s HSD, naive vs. α-synuclein overexpression, p = 0.0035, naive vs. GFP p = 0.2927 and α-syn vs. GFP p = 0.0001, from five independent replicates). (D) Representative images of naive dopaminergic neurons (top panels) and α-syn overexpressing dopaminergic neurons (bottom panels), before (left panels) and after quinpirole (10 μM) administration (right panels). (E) In naive dopaminergic neurons, bath application of quinpirole rapidly and significantly reduced the GCaMP6f fluorescence intensity (top panel) (n = 11-21, two-way ANOVA where the variables are time and treatment, p = 0.0007, from five independent biological replicates). In α-syn overexpressing neurons, bath application of quinpirole produced a smaller reduction in the peak amplitude of calcium events, as indicated by a smaller decrease in GCaMP6f fluorescence intensity (bottom panel) (n = 13-26, two-way ANOVA where the variables are time and treatment, p = 0.7259, from five independent replicates). (F) Fold change in GCaMP6f fluorescence intensity before and after drug administration (n = 11-13, one-way ANOVA followed by Tukey’s HSD, naive vs. α-syn overexpression, p = 0.0318, naive vs. GFP p = 0.8178, and α-syn vs. GFP p = 0.0057, from five independent replicates). (G) Left - A representative recording of the spontaneous firing activity of naive dopaminergic neurons before and during quinpirole (10 μM) administration exhibiting the characteristic decrease in firing activity during bath application of drug (n = 6, from three independent biological replicates). Right - A representative recording of the spontaneous firing activity of α-syn overexpressing dopaminergic neurons before and during quinpirole (10 μM) administration showing delayed and reduced reduction in firing activity (n = 8, from three independent experiments). (H) Comparison of the firing frequency of naive (black bar) and α-syn overexpressing neurons (pink bar) during quinpirole administration. The α-syn overexpressing neurons show a delayed and reduced response to quinpirole-inhibition of firing activity (n = 7 from independent experiments, 0.1812 ± SEM for naive neurons vs. 0.7930 ± SEM for α-syn treated with quinpirole, two-tailed unpaired t-test, p = 0.0356), with (I) similar interspike intervals (n = 7 from independent experiments, 11,118 ± SEM for naive neurons vs. 3786 ± SEM for α-syn neurons treated with quinpirole, twotailed unpaired t-test, p = 0.1197) and (J) firing regularity (CV of ISI - n = 7 from independent experiments, 2.06 ± SEM for naive neurons vs. 1.659 ± SEM for α-syn neurons treated with quinpirole, two-tailed t-test, p = 0.4507). The data shown in this figure are presented as mean ± SEM overlaid with data points.

While these data support the interpretation that α-syn overexpression decreases the feedback modulation of neuronal activity, they do not unequivocally show a decrease in D2 receptor activity. Dopamine interacts with multiple targets, such as dopamine transporter that also regulate neuronal excitability^46–49^ and intracellular calcium activity^46,48,50–52^. Therefore, we utilized quinpirole, a D2R-specific agonist, (10 μM) to stimulate D2R in DAT-Cre-GCaMP6f-expressing dopamine neurons containing either endogenous levels of α-syn or its overexpression (Figure 3D). Consistent with the literature^41,43,44,53^, quinpirole activation of D2 autoinhibitory receptors induced the canonical suppression of calcium dynamic in dopaminergic neurons, whereas calcium activity in α-syn overexpressing neurons did not change calcium dynamic during quinpirole administration (Figure 3E,F naive: n = 11-21, two-way ANOVA where the variables are time and treatment, p = 0.0007, from five independent replicates; α-syn: n = 13-26, two-way ANOVA where the variables are time and treatment, p = 0.7259, from five independent replicates, fold change comparisons - n = 11-13, one-way ANOVA followed by Tukey’s HSD, naive vs. α-syn, p = 0.0318, naive vs. GFP p = 0.8178, and α- syn vs. GFP p = 0.0057, from five independent replicates, Supplemental Figure 2B).

Although live cell calcium imaging provides a proxy for dopamine neuronal activity^50^, calcium imaging does not reveal changes in firing activity at the resolution of electrophysiological recordings. Therefore, as a complementary approach, we utilized whole-cell current clamp recordings to compare the spontaneous firing activity of naive and α-syn overexpressing dopaminergic neurons following quinpirole administration. Consistent with the literature, quinpirole activation of D2 autoreceptors decreased the firing activity of WT dopamine neurons^36,38,54–56^. Whereas, the response to quinpirole in α-syn overexpressing neurons is significantly attenuated (Figure 3G-J, n = 7 from independent experiments, 0.1812 ± SEM for naive neurons vs. 0.7930 ± SEM for α-syn treated with quinpirole, two-tailed unpaired t-test, p = 0.0356, interspike interval - n = 7 from independent experiments, 11,118 ± SEM for naive neurons vs. 3786 ± SEM for α- syn neurons treated with quinpirole, two-tailed unpaired t-test, p = 0.1197, firing regularity (CV of ISI) - n = 7 from independent experiments, 2.06 ± SEM for WT neurons vs. 1.659 ± SEM for α-syn neurons treated with quinpirole, two-tailed t-test, p = 0.4507). Surprisingly, we observed that the longer the exposure to quinpirole, the firing activity of α-syn overexpressing dopamine neurons began to resemble naive dopaminergic neurons at baseline (before drug application, end of quinpirole treatment of α-syn neuron recording, Figure 3G - right, pink). Collectively, these data support the idea that α-syn overexpression reduces the efficacy of D2 receptor autoinhibition in dopamine neurons and a prolonged activation of D2 receptor can potentially restore this deficit.

### α-synuclein overexpression increases intra- and extracellular dopamine levels and tyrosine hydroxylase expression

Our findings so far suggest that α-syn may induce a feed-forward adaptive mechanism that decreases the ability of inhibitory D2 autoreceptors to act as a break on neuronal excitability and increasing extracellular dopamine levels^33,36^. To test this hypothesis, we used two complementary approaches to measure intra- and extracellular dopamine levels. First, we used GRAB_DA4.4_ (G protein-coupled receptor [GPCR]-activation-based DA) sensor-expressing HEK293 cells to measure extracellular dopamine levels. GRAB_DA4.4_ is a genetically encoded fluorescent dopamine sensor, engineered by coupling a conformationally-sensitive circular-permutated EGFP (cpEGFP) to D_2_ receptor. In GRAB_DA4.4-_expressing HEK293 cells, dopamine binding to the sensor induces a conformational change which results in a robust increase in fluorescence signal in a concentration-dependent manner (Figure 4A-B). Constitutive GRAB_DA4.4_ fluorescence signal in the absence of dopamine neurons (in the culture) was obtained at the beginning of each experiment, where the cells were plated in similar conditions sans neurons (F_c_, Figure 4C). To compare baseline dopamine release amongst the experimental groups, the average ratio of fluorescence signal of cells adjacent to the soma and neuronal processes to the average ratio of fluorescence signal of GRAB_DA4.4_ cells (only) were calculated (F_baseline_ = (F_GRABDA4.4 cells grown with neurons_ - F_c_) *I* F_c_). To compare KCl stimulation of dopamine release, the changes on the average fluorescence signal of cells adjacent to the soma and neuronal processes before and after KCl were calculated (ΔF/F = (F_stimulated_ - F_baseline_) *I* F_baseline_). Co-culture of GRAB_DA4.4_ cells with dopamine neurons 20-24 hours prior to live cell confocal imaging enabled realtime detection of endogenous dopamine released from the neurons at baseline, i.e. spontaneous dopamine release (Figure 4D, E). Using a double-blinded experimental design, we found a significantly higher basal extracellular dopamine level, as measured by a higher GRAB_DA4.4_ fluorescence signal around the soma and neuronal processes of α-syn overexpressing neurons (Figure 4F, n = 10 from three independent replicates; the data are means ± SEM, one-way ANOVA followed by Tukey’s HSD, F(2,167) = 13.05, naive vs. α-syn p = 0.0007, WT vs. GFP p = 0.9866 and GFP vs. α-syn p = 0.0001). These data support the interpretation that α-syn-induced stimulation of spontaneous neuronal activity leads to increased extracellular dopamine levels. As a positive control group, we measured the GRAB_DA4.4_ fluorescent signal around the soma and neuronal processes following KCl (90mM) stimulation of dopamine release^57^. KCl-induced neuronal depolarization^58,59^ produced a robust fluorescence increase that was not different amongst the experimental groups (Figure 4G, n = 10 from three independent replicates; the data are mean ± SEM analyzed by one-way ANOVA followed by Tukey’s HSD, F(2,68) = 0.2540, naive vs. α-syn p = 0.9872, naive vs. GFP p = 0.8651 and GFP vs. α-syn p = 0.7827). Consistent with our previous report, these data suggest that α-syn overexpression decreases dopamine uptake via the dopamine transporter (DAT)^60^ and increases the DAT-mediated dopamine efflux^61^ leading to increased extracellular dopamine levels. Collectively, these data provide a reasonable cellular mechanism for the puzzling observation by Lam et al., that in mice overexpressing α-syn there is an initial increase in extracellular dopamine levels in the striatum prior to neuronal death^62^.

**Figure 4.**
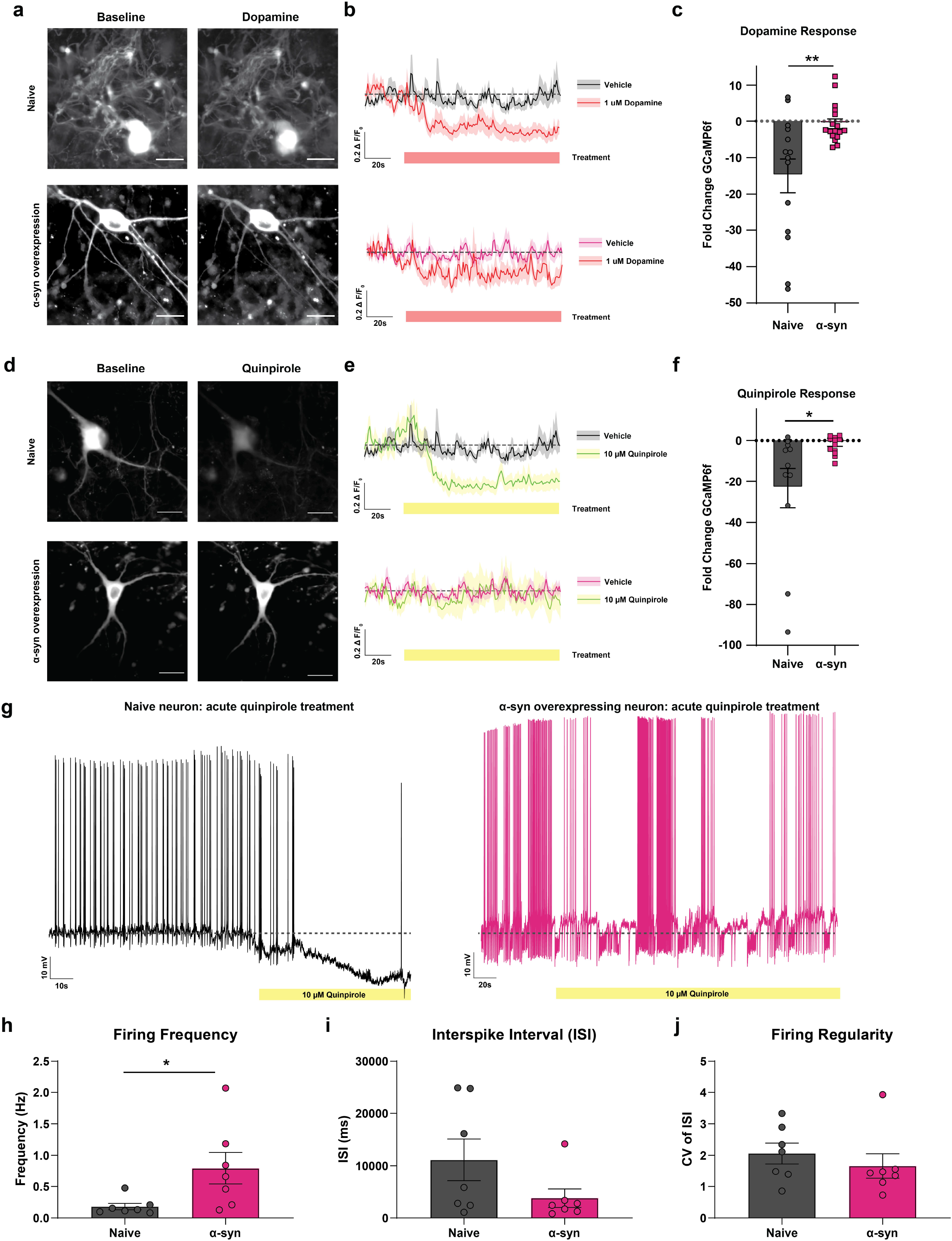
Overexpression of α-synuclein increases intra- and extracellular dopamine levels with concurrent increased tyrosine hydroxylase expression. (A) Schematic and representative baseline-subtracted images of GRAB_DA4.4-_expressing HEK293 (GRAB_DA4.4-_HEK) cells exposed to increasing concentration of dopamine in the imaging solution. Scale bar = 20 μm. (B) A standard curve was generated by plotting the fluorescence intensity of GRAB_DA4.4-_HEK cells against known extracellular dopamine concentrations (R^2^=0.98). (C) Constitutive GRAB_DA4.4-_HEK cell fluorescence signal in absence of dopamine neurons (in culture) was obtained at the beginning of each experiment, where the cells were plated in similar conditions sans neurons (F_c_). Scale bar = 10 μm. (D) Schematic of experimental procedure where GRAB_DA4.4-_HEK cells were seeded into dopaminergic cultures. In the presence of dopamine, GRAB_DA4.4-_HEK cells rapidly increase in fluorescence intensity. GRAB_DA4.4-_HEK cells were co-cultured for 20-24 hours with DATCre-GCaMP6f midbrain dopamine neurons prior to experimentation. Viral transduction of floxed tdTomato (red) enabled isolation of the GCaMP6f signal from GRAB_DA4.4_ signal. (E) Baseline fluorescence levels denote the basal, unstimulated, and spontaneous dopamine release from the neurons. To compare baseline dopamine release amongst the experimental groups, the average ratio of the fluorescence signal of the cells adjacent to neuron soma and neuronal processes to the average ratio of GRAB_DA4.4-_HEK cells (only) were calculated (Relative Fluorescence = (FGRAB_DA4.4-_HEK cells grown with neurons - F_c_) /F_c_). To compare KCl-stimulated dopamine release, the changes on the average fluorescence signal of cells adjacent to the neuron soma and neuronal processes before and after KCl were calculated (Relative Fluorescence = (F_stimulated_ - F_baseline_) /F_baseline_). GRAB_DA4.4-_HEK cells co-cultured with α-syn overexpressing neurons show significantly higher basal extracellular dopamine release (relative fluorescence) compared to naive and AAV-TH-GFP expressing neurons. Scale bar = 50 μm. (F) Bar graph shows GRAB_DA4.4-_HEK cells co-cultured with α-syn overexpressing neurons show higher basal fluorescence, indicating higher baseline dopamine release (n = 10 from three independent replicates; the data are means ± S.E.M., one-way ANOVA followed by Tukey’s HSD, F(2,167) = 13.05, naive vs. α-syn p = 0.0007, naive vs. GFP p = 0.9866 and GFP vs. α-syn p = 0.0001). (G) Bath application of 90 mM KCl (positive control for maximal dopamine release) reveals similar relative fluorescence responses across groups, indicating similar maximal dopamine release (n = 10 from three independent replicates; the data are mean ± S.E.M. analyzed by one-way ANOVA followed by Tukey’s HSD, F(2,68) = 0.2540, naive vs. α-syn p = 0.9872, naive vs. GFP p = 0.8651 and GFP vs. α-syn p = 0.7827). (H, I) To further directly measure intra- and extracellular dopamine levels, HPLC analyses was used to complement the GRAB_DA4.4-_ HEK results. HPLC quantification of dopamine levels in the in the media (H, extracellular milieu) and cell lysate (I, intracellular milieu) revealed increased intracellular and extracellular dopamine levels in α-syn overexpressing neurons compared to naive and AAV-TH-GFP-expressing neurons (n = 8 each, from 8 independent replicates, one-way ANOVA followed by Tukey’s HSD, intracellular: F(3,22) = 9.376, naive vs. α-syn p = 0.0059, naive vs. GFP p = 0.5164 and GFP vs. α-syn p = 0.0003; and extracellular: F(3,22) = 9.525, naive vs. α-syn p = 0.01, naive vs. GFP p = 0.5235 and GFP vs. α-syn p = 0.0005). (J) Schematic diagram of quantitative ELISA experimental design for TH in dopaminergic neurons. (K) Standard curve for TH sandwich ELISA shows average absorbance values for each purified TH protein concentration from multiple consecutive experiments (R^2^=0.99). (L) TH protein levels were detected and quantified in positive control groups, PC12 cells, whereas, no protein was detected in the negative control group, HEK293 cells. (M) α-Syn overexpressing neurons exhibited increased levels of TH compared to naive and AAV-TH-GFP controls (n = 8-10, one-way ANOVA followed by Tukey’s HSD, F(2,23) = 4.488, naive vs. α-syn p = 0.031, naive vs. GFP p = 0.7983, and GFP vs. α-syn p = 0.0491). These experiments were performed through a double-blinded experimental design.

As a complementary approach, via a double-blinded experimental design, we used HPLC to measure dopamine level in the external milieu of naive dopaminergic neurons and dopaminergic neurons overexpressing α-syn. HPLC analysis (see Methods) showed a significantly higher extracellular dopamine in α-syn overexpressing neurons compared to naive neurons or dopaminergic neurons expressing GFP (Figure 4H, I; n = 8 each, from 8 independent replicates; one-way ANOVA followed by Tukey’s HSD, F(3,22) = 9.525, WT vs. α-syn p = 0.01, WT vs. GFP p = 0.5235 and GFP vs. α-syn p = 0.0005). Collectively, these data, combined with live cell detection of extracellular dopamine levels, support the notion that α-syn modulation of dopaminergic neuronal activity leads to increased extracellular dopamine that could be due to increased neuronal activity, increased dopamine synthesis, or both possibilities. Since we have already examined the former (Figure 4F,H), to test the latter possibility, we used HPLC to measure intracellular dopamine levels via a double-blinded experimental design (described in Methods). The measurement of dopamine in the cell lysate of naive and α-syn overexpressing neurons revealed a significantly higher intracellular dopamine (Figure 4 I, n = 8 each, from 8 independent replicates, one-way ANOVA followed by Tukey’s HSD, F(3,22) = 9.376, naive vs. α-syn p = 0.0059, naïve vs. GFP p = 0.5164 and GFP vs. α-syn p = 0.0003). These data suggest that the decreased autoinhibition of dopamine neurons following α-syn overexpression not only increases neuronal excitability, but also dysregulates dopamine synthesis and secretion. Furthermore, in Figure 2, we showed that increased neuronal α-syn increases the magnitude and duration of intracellular calcium burden, which would promote increased basal dopamine release.

Multiple mechanisms likely contribute to the increased intracellular dopamine following α-syn overexpression. For example, increased dopamine uptake via the dopamine transporter (DAT), decreased DAT-mediated dopamine efflux, increased expression of tyrosine hydroxylase (TH), a key enzyme involved in dopamine synthesis, or a combination of these mechanisms would possibly contribute to a higher intracellular dopamine level. Previously, we and others have shown that α-syn overexpression reduces dopamine recycling by reducing dopamine uptake^60,63,64^. In addition we have shown that α-syn overexpression, increases reverse transport of dopamine i.e. dopamine efflux^61^, without changing surface DAT levels. Therefore, α-syn regulation of dopamine uptake or dopamine efflux would decrease intracellular dopamine, not increase it.

Whereas α-syn regulation of DAT activity predicts a decrease of intracellular dopamine level^60,61,63–65^, D2 receptor activity negatively regulates TH protein levels as a compensatory mechanism to down-regulate dopamine synthesis^36,43,66–69^. As shown in Figure 2 and Figure 3, we found a reduction in the canonical D2 receptor autoinhibition of dopamine neurons, likely modulating downstream signaling cascades that can regulate TH protein levels. Therefore, next we tested the hypothesis that increased TH levels caused by reduced D2 autoinhibition in α-syn overexpressing neurons (as observed in Figures 2, 3) underlies the increased intracellular dopamine^66,69–74^. Since the frequently used approaches of western blotting or immunocytochemistry do not provide purely quantitative data of protein expression to test this hypothesis, we developed an ELISA to quantify TH levels in α-syn overexpressing neurons (Figure 4 J). For these experiments we used HEK293 cells as a negative control group and PC12 cells as a positive control group (Figure 4 L). Purified full-length recombinant TH protein was used to generate a standard curve (Fig 4 K). α-syn overexpressing neurons show significantly higher TH levels, compared to naive and control conditions (Figure 4M, n = 8-10, one-way ANOVA followed by Tukey’s HSD, F(2,23) = 4.488, naive vs. α-syn p = 0.031, naive vs. GFP p = 0.7983, and GFP vs. α-syn p = 0.0491). While ELISA provides quantitative data for total TH level across these experimental groups, a limitation of this assay is that it cannot discriminate TH phosphorylation that is associated with TH activity and thus dopamine synthesis^66,75,76^. Nevertheless, these data support the interpretation that increased intracellular dopamine in α-syn overexpressing neurons, at least in part, is due to increased TH protein levels.

### Altered neural dynamics mediated by α-synuclein may emerge from altered D2 activity and expression patterns

Our data, so far, support the interpretation that the canonical D2 receptor autoinhibitions, such as inhibitory modulation of spontaneous firing activity, is reduced in α-syn overexpressing neurons. While D2 receptor agonist quinpirole silenced naive dopamine neurons, the response to quinpirole in α-syn overexpressing neurons is significantly delayed and reduced, possibly due to desensitization or reduced activity of the D2 receptor (Figure 3). Therefore, next we tested the hypothesis that blockade of D2 receptor in naive dopaminergic neurons simulates the firing activity observed in α-syn overexpressing neurons. We performed whole-cell current clamp recordings to measure spontaneous firing activity of dopaminergic neurons before and during bath application of sulpiride (D2 antagonist, 5 μM). In naive dopaminergic neurons, bath application of sulpiride produced burst firing patterns with intermediated periods of quiescence and firing frequencies resonant with neuronal activity seen in α-syn overexpressing dopaminergic neurons (Fig 5 A-E, n = 8 from 3 independent biological replicates, two-tailed unpaired t-test, firing frequency: 1.766 ± SEM naive vs. 2.805 ± SEM α-syn overexpressing neurons, p = 0.148, ISI - 1791 ± SEM naive vs. 1221 ± SEM α-syn overexpressing neurons p = 0.1147, CV of ISI - 1.642 naive ± SEM vs. 1.412 ± SEM α-syn overexpressing neurons, p = 0.456). These data support the hypothesis that in α-syn overexpressing dopamine neurons, the reduced functional availability of D2-mediated response could be due to receptor desensitization^77,78^, decreased membrane expression of D2 receptors, or a combination of these possibilities.

**Figure 5.**
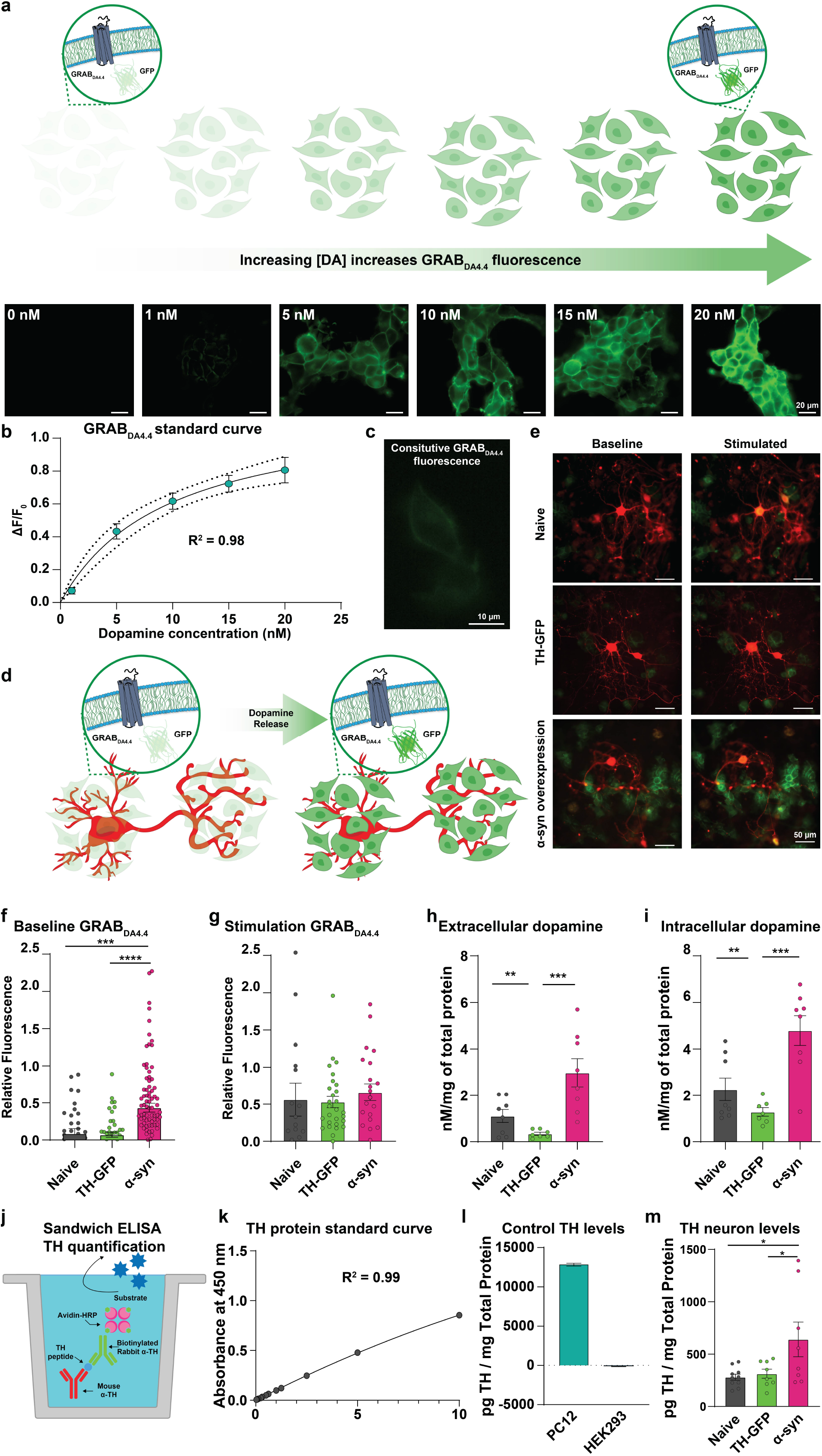
D2 receptor antagonism in dopaminergic neurons mimics burst firing pattern with a significantly higher firing frequency observed in α-synuclein overexpressing dopamine neurons that presents with lower membrane/cytoplasmic D2 ratio. (A-B) Representative whole cell current-clamp recordings of spontaneously active naive (A, black) and α-syn overexpressing (B, pink) dopaminergic neurons during sulpiride (D2 antagonist, 5μM) bath application. (C-E) Bar graph shows firing frequency (C), interspike interval (ISI) (D), and firing regularity (E) during bath application of sulpiride (5μM), revealing D2 antagonism in naive dopaminergic neurons promotes firing rates, interspike intervals, and regularity comparable to neurons overexpressing α-syn (n = 8 from 3 independent biological replicates, two-tailed unpaired t-test, firing frequency: 1.766 ± SEM naive vs. 2.805 ± SEM α-syn overexpressing neurons, p = 0.148, ISI - 1791 ± SEM naive vs. 1221 ± SEM α-syn overexpressing neurons p = 0.1147, CV of ISI - 1.642 naive ± SEM vs. 1.412 ± SEM α-syn overexpressing neurons, p = 0.456). (F-G) Membrane biotinylation experiments. Membrane and cytoplasmic D2 factions were isolated from a separate set of naive and α-syn overexpressing neurons, prepared identically, via a double blinded experimental design. (F) ?-syn overexpressing neurons exhibit an increased trend in levels of cytoplasmic D2 receptor relative to total D2 receptor levels (n = 3 from independent replicates, two-tailed unpaired t-test p = 0.1115). (G) The ratio of average membrane D2R to the average cytoplasmic D2R significantly decreases in α-syn overexpressing neurons (n = 3 from independent replicates, two-tailed unpaired t-test p = 0.0178). The data are presented as mean ± SEM with independent replicate data points overlaid.

To investigate if α-syn overexpression in dopaminergic neurons alters D2 receptor expression, we performed cell surface biotinylation of D2 receptor via a double-blinded experimental design. We found increased ratio of average cytoplasmic D2R to average total D2R in α-syn overexpressing neurons (Figure 5 F, n = 3 from independent replicates, two-tailed unpaired t-test, p = 0.1115). Whereas the ratio of average membrane D2R to the average cytoplasmic D2R decreased compared to naïve neurons (Figure 5 G, from 3 independent replicates, two-tailed unpaired t-test, p = 0.0178). The double immunocytochemistry of fixed but not permeabilized dopamine neurons stained for both D2R and an integral membrane protein such as Na+IK+-ATPase, or GM1-CTxB would have been a suitable complementary approach to examine membrane localized D2R across the experimental groups in this study. However, the frequently used D2R antibodies in the field^77–80^ are raised against the intracellular N-terminal domain of the receptor. This limitation decreases the confidence in identification of membrane vs. intracellular protein levels. Similar technical limitation applies to the single cell qPCR assay, where total transcript levels do not necessarily reflect functional D2Rs at the membrane. The latter limitation somewhat applies to the biotinylation assay used in this study. Unless an antibody is raised against the active or inactive form of the receptor, a biotinylation assay detects both functional and desensitized receptors. Therefore, although our data suggest membrane D2Rs are decreased in α-syn overexpressing neurons, it is possible that the detected membrane D2Rs are desensitized, i.e. a lesser receptor-effector coupling^81–84^. Therefore, live cell functional assays, such as electrophysiology and calcium imaging, combined with pharmacological manipulations are more reliable strategies to assess the mechanism of α-syn regulation of neuronal activity.

### α-synuclein overexpression reduces arborization of dopamine neurons and pretreatment with a D2 receptor agonist partially rescues the detrimental impact of α-synuclein

Dopaminergic neurons have extensive axonal arborizations and large terminal fields^85–87^, where one dopamine neuron is estimated to have ~245,000 release sites^88,89^. Studies in animal models of PD and postmortem data in human PD^90^ show decreased axonal complexity and dendritic arborization, reduction of number of axon terminals and global neuronal size precede neuronal death^63,88,91^. Our data suggest that prior to cell death, via a D2 receptor mechanism, α-syn overexpression can induce neuronal disinhibition, leading to increased intra- and extracellular dopamine levels that are implicated in increased neuronal vulnerability^13,92,93^. Therefore, we investigated the potential link between α-syn-mediated dopamine neuronal dysfunction and neuronal complexity.

Sholl analysis entails using concentric circles around the soma of a neuron, with dendritic fields intersecting these concentric circles counted as a measure of differences in neuronal complexity (Figure 6 A-D). This analytical approach estimates^87,94–96^ neuronal complexity via assessment of projection area, number of intersections as a measure of neuronal arborization, projection field perimeter, neuronal arborization width, and circularity of dendritic arborization^94,96–100^. Compared to naive dopaminergic neurons, α-syn overexpressing neurons exhibit a lower degree of neuronal arborization (Figure 6 E, F, H), reduced projection arborization area (Fig. 6 J), reduced circularity of dendritic arborization (Fig 6 K), smaller dendritic projection perimeter (Fig. 6 L), and smaller arborization width (Fig. 6 M), but no change in the soma area (Fig 6 I, one-way ANOVA followed by Tukey’s test, naive n = 190, α-syn n = 114, intersections: naive vs. α-syn p = 0.0009, circularity: naive vs. α-syn p = 0.0021, outer perimeter: naive vs. α-syn p = 0.0001; width: WT vs. α-syn p = 0.0001, projection area: naive vs. α-syn p = 0.0001, soma area: naive vs. α-syn p = 0.67, from at least three independent biological replicates). The loss of neuronal complexity and decreased dendritic arborization found in this study are consistent with morphological data in postmortem PD samples^90^, potentially informing the progression of α-syn-induced pathology prior to neuronal loss.

**Figure 6.**
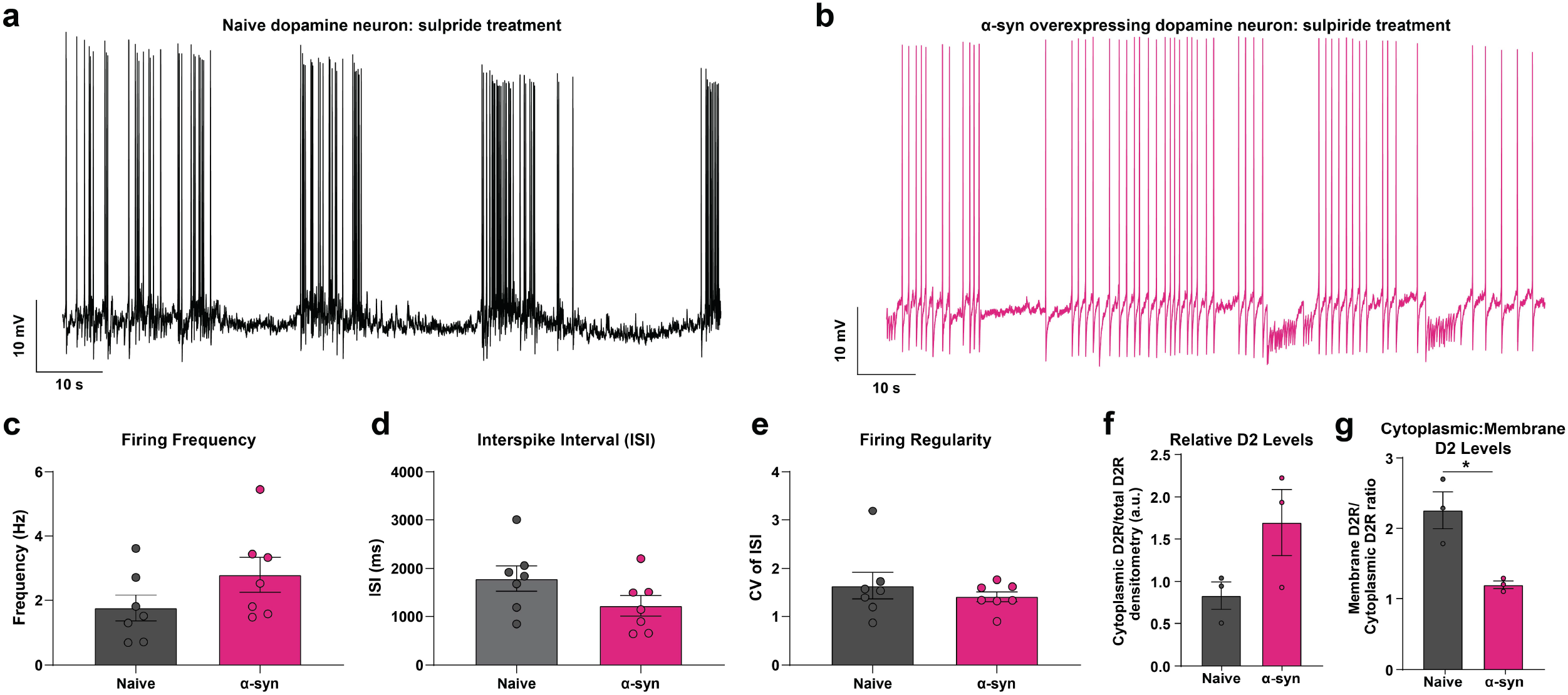
α-Synuclein overexpression reduces arborization of dopaminergic neurons and treatment with a D2 receptor agonist partially rescues the detrimental impact of α-synuclein. (A) Schematic image explaining the parameters used in the morphological examination (B-D) Representative binarized images of naive (B), α-syn overexpressing (C), and quinpirole pretreated α-syn overexpressing neurons (D). (E-H) Sholl intersection profiles of untreated naive (E), untreated α-syn-overexpressing (F), and quinpirole pretreated α-syn-overexpressing neurons (G) followed by measurement of area under curve (H). Significantly fewer intersections in α-syn overexpressing neurons were observed compared to naive neurons, indicating a marked reduction in neuronal arborization (Tukey’s HSD following one-way ANOVA, naive n = 190, α-synuclein, n = 114, naive vs. α-synuclein, p = 0.0009, from at least three independent replicates). Sholl analyses revealed that 48 hours of quinpirole (0.5 μM) pretreatment induces a partial restoration of neural arborization complexity (described above) compared to untreated α-syn overexpressing neurons (Tukey’s HSD following one-way ANOVA, naive n = 190, α-syn n = 114 and α-syn treated with quinpirole n = 32, naive vs. α-syn p = 0.0009, naive vs. α-syn treated with quinpirole p = 0.5131, and α-syn vs. α-syn treated with quinpirole p = 0.0041, from at least three independent replicates). (I) Somatic areas were found to be comparable between experimental groups (Tukey’s HSD following one-way ANOVA, naive n = 190, α-syn n = 114, naive vs. α-syn p = 0.67, from at least three independent biological replicates). (J) α-Syn overexpressing dopaminergic neurons project over a much smaller area than naive neurons (Tukey’s HSD following one-way ANOVA, naive n = 190, α-syn n = 114, naive vs. α-syn p = 0.0001, from at least three independent biological replicates). (K-M) Morphological measures show detrimental morphological changes in α-syn overexpressing neurons compared to naive control neurons (Tukey’s HSD following one-way ANOVA, naive n = 190, α-syn n = 114, circularity: naive vs. α-syn p = 0.0021, outer perimeter: naive vs. α-syn p = 0.0001; width: naive vs. α-syn p = 0.0001, from at least three independent replicates). (K) Circularity of the neurons was measured as ratio of projection area and the projection perimeter ((4·π·area)/perimeter^2^). (L) Projection field perimeter was defined as the perimeter of the neuronal projection field at the outermost process. (M) Neuronal arborization width was defined as the ratio of projection area to the longest dendritic length of each neuron. D2 receptor agonist partially rescues the detrimental impact of α-synuclein. Quinpirole treatment of α-syn overexpressing neurons rescued changes in (J) projection area (Tukey’s HSD following one-way ANOVA, naive n = 190, α-syn n = 114 and α-syn treated with quinpirole n = 32, naive vs. α-syn p = 0.0001, naive vs. α-syn treated with quinpirole p = 0.1136, and α-syn vs. α-syn treated with quinpirole p = 0.0486, from at least three independent replicates), (K) neuronal circularity, (L) projection field perimeter, and (M) arborization width (Tukey’s HSD following one-way ANOVA (put in the F(df1,df2) here), naive n = 190, α-syn n = 114, and α-syn treated with quinpirole n = 32, circularity: naive vs. α-syn p = 0.0021, naive vs. α-syn treated with quinpirole p = 0.4452, and α-syn vs. α-syn treated with quinpirole p = 0.0046; outer perimeter: naive vs. α-syn p = 0.0001, naive vs. α-syn treated with quinpirole p = 0.2250, and α-syn vs. α-syn treated with quinpirole p = 0.0312; width: naive vs. α-syn p = 0.0001, naive vs. α-syn treated with quinpirole p = 0.1419, and α-syn vs. α-syn treated with quinpirole p = 0.0187, from at least three independent biological replicates). (N) Dopaminergic neuron counts revealed α-syn overexpression decreases neuronal survival, which is rescued when pretreated with quinpirole (0.5 μM for 48 hours) (Tukey’s HSD following one-way ANOVA, naive vs. α-syn p = 0.0008, naive vs. α-syn treated with quinpirole p = 0.1364, and α-syn vs. α-syn treated with quinpirole p = 0.0021, from two independent replicates).

The unexpected observation, that prolonged bath application of D2 receptor agonist began to restore the firing properties of α-syn overexpressing neurons to the levels measured in WT dopamine neurons at baseline (end of quinpirole treatment of α-syn neuron recording, Figure 3G - right, pink), suggests that pharmacological activation of D2 receptor is a potential target to alleviate α-syn pathology prior to neuronal death. This hypothesis is consistent with reports showing that D2 receptor activation sustains structural plasticity of dopaminergic neurons by maintaining their dendritic arborization^93^,^101–104^. Therefore, we tested whether the “retrograde axonal degeneration” hypothesis described in the literature^105^,^106^ could be ameliorated through D2 receptor activation, leading to restoration or reduction of the declining degree of neuronal complexity^93^,^107^,^108^. We measured neuronal complexity when α-syn overexpressing neurons were pretreated with quinpirole (0.5 μM) for 48-hours. Surprisingly, we found that quinpirole produced a marked improvement in dendritic arborization of α-syn overexpressing neurons (Figure 6D, G). Detailed morphometrical analysis revealed no change in somatic size of neurons across all experimental groups; whereas, quinpirole restored neuronal circularity, perimeter of the projection field, projection perimeter, and width of α-syn overexpressing dopamine neurons, to the level measured in naive untreated neurons shown in Figure 6 B-M (naive n = 190, α-syn n = 114 and α-syn + quinpirole n = 32, from at least three independent replicates; circularity: naive vs. α-syn p = 0.0021, naive vs. α-syn + quinpirole p = 0.4452, and α-syn vs. α-syn + quinpirole p = 0.0046; outer perimeter: WT vs. α-syn p = 0.0001, WT vs. α-syn + quinpirole p = 0.2250, and α-syn vs. α-syn + quinpirole p = 0.0312; shape factor: naive vs. α-syn p = 0.0021, naive vs. α-syn + quinpirole p = 0.5574, and α-syn vs. α-syn + quinpirole p = 0.0086; width: naive vs. α-syn p = 0.0001, WT vs. α-syn + quinpirole p = 0.1419, and α-syn vs. α-syn + quinpirole p = 0.0187, one-way ANOVA followed by Tukey’s HSD).

Next, we examined the impact of α-syn overexpression on neuronal survival by counting TH-positive neurons via immunocytochemistry. Compared to naive neurons, we found significantly fewer TH positive neurons following α-syn overexpression; quinpirole-mediated activation of D2 receptors (0.5 μM for 48 hours) prevented the neuronal loss (Fig 6 N: WT vs. α-syn p = 0.0008, WT vs. α-syn + quinpirole p = 0.1364, and α-syn vs. α-syn + quinpirole p = 0.0021, from three independent biological replicates, one-way ANOVA followed by Tukey’s HSD). The increased neuronal survival following quinpirole pretreatment is consistent with previous reports^93^,^101–104^ and support the interpretation that there is a correlation between α-syn modulation of neuronal complexity, neuronal vulnerability, and neuronal loss^109–111^. Neuronal survival is not equivalent to neuronal viability. While these data suggest pretreatment with a D2 receptor agonist increases neuronal survival following α-syn overexpression, they do not demonstrate a restoration of neuronal activity. Therefore, next, we asked whether quinpirole pretreatment prevents the perpetual increase in action potential frequency, increased intra- and extracellular dopamine levels, and elevated intraneuronal calcium dynamics in α-syn overexpressing neurons.

### D2 receptor activation partially restores neuronal activity in α-synuclein-overexpressing dopamine neurons

The energy homeostasis principle suggests, the balance between energy income, expenditure, and availability are the key parameters in determining neuronal endurance^112^. Action potentials impose the highest energy demands on neurons^88,112,113^. In addition, dopamine metabolism is strongly linked to oxidative stress, as its degradation generates reactive oxygen species^13,114,115^ that have shown to increase the vulnerability of dopamine neurons to oxidative stress^87,115–121^. So far, we have identified multiple interrelated mechanisms that would add fuel to vulnerability of α-syn overexpressing dopamine neurons. We identified a perpetual increase in action potential frequency, increased intra- and extracellular dopamine levels, and elevated intraneuronal calcium dynamics in α-syn overexpressing neurons, that are directly or indirectly related to decreased D2R activity. The unexpected observation that bath application of D2R agonist increased neuronal survival and nearly restored neuronal complexity of α-syn overexpressing neurons to the levels measured in naive dopaminergic neurons at baseline, suggests pharmacological activation of D2 receptors might be a possible target to alleviate the untoward consequences of α-syn overexpression on neuronal activity prior to neuronal death. To test this hypothesis, we treated α-syn overexpressing neurons with 0.5 μM quinpirole for 48 hours before assessing calcium dynamics, spontaneous firing activity, and dopamine release and synthesis in these neurons (Figure 7). This data is compared to the previous data which is overlaid with a black dotted line that represents the average values measured for untreated naive neurons and the pink dotted line represents the average values for untreated α-syn overexpressing neurons. We found that a prolonged D2R activation partially restores calcium dynamics in these neurons, approximating calcium dynamics measured in untreated naive neurons (Figure 7 A-F, n = 28 quinpirole treated α-syn overexpressing neurons, two-tailed unpaired t-test, α-syn vs. α-syn pretreated with quinpirole p = 0.2024 event rate, p = 0.0277 event widths, p = 0.6204 event height), suggesting a restoration of calcium homeostasis in these neurons that might be causal or a consequence of a shift in neuronal activity. To test this hypothesis, we employed whole-cell current clamp recordings to measure the spontaneous firing activity of α-syn overexpressing neurons after treatment with quinpirole (0.5 μM for 48 hours). Quinpirole pretreatment on α-syn overexpressing neurons decreased the burst firing frequency, shortened the intermediated periods of quiescence and restored firing regularity, near the values measured in untreated naive neuron’s firing pattern (Figure 7G-K, n = 7 from three independent biological replicates, 1.325 ± SEM for quinpirole-treated α-syn overexpressing neurons, two-tailed unpaired t-test, α-syn vs. α-syn pretreated with quinpirole p = 0.0342 for firing frequency, p = 0.1053 for ISI, p = 0.4778 for CV of ISI). These results suggest that dysregulation of D2 receptor in α-syn overexpressing dopamine neurons can be partially rescued with prolonged activation of the remaining functional D2 receptors on the cell surface

**Figure 7.**
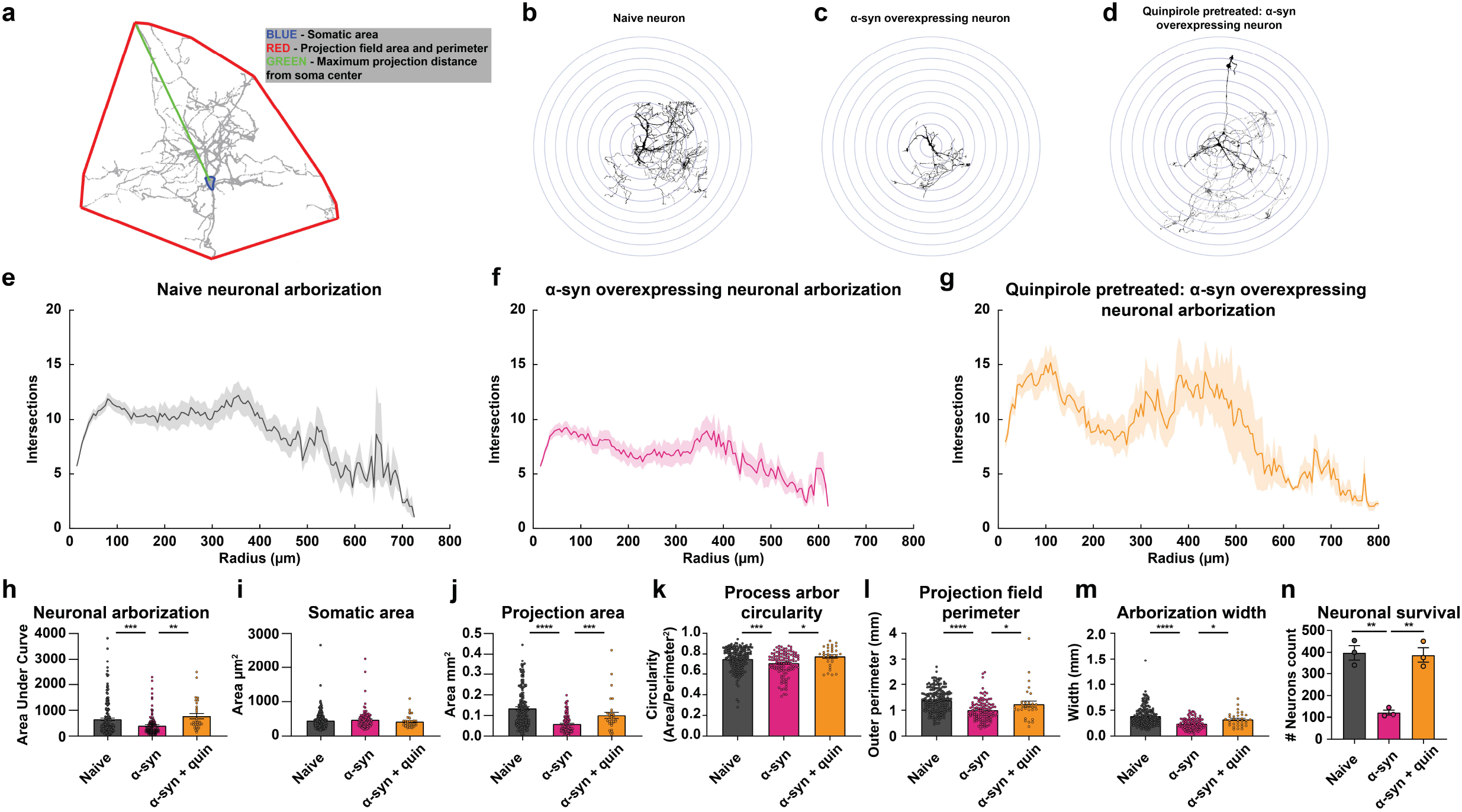
Pretreatment with D2 receptor stimulation partially restores neuronal activity in α-synuclein overexpressing dopamine neurons. (A) To test whether prolonged D2 receptor stimulation ameliorates the impact of α-syn overexpression, dopamine neuronal cultures were incubated with a D2 receptor agonist (quinpirole, 0.5 μM for 48 hours). (B) Spontaneous GCaMP6f calcium activity in a dopaminergic neuron overexpressing α-syn pretreated with quinpirole illustrating a partial restoration of calcium dynamics similar to untreated naive neurons. (C) Calcium events in B were identified as the fluorescent signal increases at least two standard deviations above the fluorescence baseline value. (D-F) Event rate (D), event width (E), and event amplitude (F) after quinpirole pretreatment. Quinpirole pretreatment reverts the α-syn modulation of calcium homeostasis by preventing the event width broadening, but not the increasing event amplitude or changing the event rates (n = 28 quinpirole treated α-syn overexpressing neurons, two-tailed unpaired t-test, α-syn vs. α-syn pretreated with quinpirole p = 0.2024 event rate, p = 0.0277 event widths, p = 0.6204 event height). Throughout this figure previous data is overlaid with a black dotted line that represents the average values measured for untreated naive neurons and a pink dotted line represents the average values for untreated α-syn overexpressing neurons. Representative of firing activity of an untreated naive neuron (G) and quinpirole pretreated α-syn overexpressing neuron (H). (I-K) Quinpirole treatment partially restores the firing activity of α-syn overexpressing neurons. Quantification of firing frequency (I), interspike interval (J), and firing regularity (K) in quinpirole pretreated α-syn dopamine neuron (n = 7 from three independent biological replicates, 1.325 Hz ± SEM for quinpirole-treated α-syn overexpressing neurons, two-tailed unpaired t-test, α-syn vs. α-syn pretreated with quinpirole p = 0.0342 for firing frequency, p = 0.1053 for ISI, p = 0.4778 for CV of ISI). (L) 20-24 hours prior to experimentation, GRAB_DA4.4_ expressing HEK293 (GRAB_DA4.4-_HEK) cells were seeded at untreated naive (left), untreated α-syn overexpressing (middle) and quinpirole pretreated α-syn overexpressing (right) midbrain neuronal cultures (dopaminergic neurons identified through floxed expression of tdTomato). (M) Quinpirole pretreatment rescued increased dopamine level (quantified as the relative GRAB_DA4.4_ fluorescence levels) in α-syn-overexpressing neurons to level similar to untreated naive neurons (n = 6 from three independent biological replicates; data shown as mean ± SEM, one-way ANOVA followed by Tukey’s HSD, naive vs. α-syn treated with quinpirole p = 0.9948, α-syn vs. α-syn treated with quinpirole p = 0.0003) (N,O) GRAB_DA4.4_ signals provide qualitative representation of α-syn-induced dopamine release. Therefore, to corroborate these data, we performed HPLC to quantify intracellular (lysate) and extracellular (media) dopamine levels. The HPLC quantification of dopamine levels revealed that quinpirole pretreatment of α-syn overexpressing neurons reduces both extracellular (N) and intracellular (O) dopamine levels compared to untreated α-syn overexpressing neurons (n = 3 each, from 3 independent biological replicates, one-way ANOVA followed by Tukey’s HSD, intracellular: α-syn vs. α-syn treated with quinpirole p = 0.0325 and naive vs. α-syn treated with quinpirole p = 0.9959; extracellular: α-syn vs. α-syn treated with quinpirole p = 0.0449 and naive vs. α-syn treated with quinpirole p = 0.6197). (P) Quantification of TH expression by ELISA. Compared to untreated α-syn overexpressing neurons, quinpirole pretreatment decreased TH protein levels in α-syn overexpressing neurons (n = 3, oneway ANOVA followed by Tukey’s HSD, naive vs. α-syn treated with quinpirole p = 0.4288 and α-syn vs. α-syn treated with quinpirole p = 0.6809). The experiments were performed via double-blinded experimental design. Data are presented as mean ± SEM with data points shown.

The observed changes in neuronal responses and calcium activity following extended D2 receptor activation could be predictive of downstream changes in dopamine synthesis in α-syn overexpressing neurons. To test the hypothesis that D2 receptor activation decreases α-syn modulation of dopamine release, we measured extracellular dopamine levels via two complementary approaches: live-cell imaging and HPLC. To measure D2R-mediated modulation of baseline extracellular dopamine levels (0.5 μM quinpirole 48 hours) in α-syn overexpressing neurons, we co-cultured GRAB_DA4.4_-expressing cells with the quinpirole-treated, α-syn overexpressing dopamine neurons for 20-24 hours prior to imaging. Quinpirole pretreatment of α-syn overexpressing neurons decreased basal GRAB_DA4.4_ fluorescence (used as a proxy to measure basal dopamine release) around the soma and dendritic fields (Figure 7 L, M), comparable to values measured in naive untreated neurons shown in Figure 4 (black dotted line in Figure 7). Consistent with the literature^36,54,122–125^, quinpirole silenced naive dopamine neurons and decreased basal extracellular dopamine (n = 6 from three independent replicates; data shown as ±SEM, one-way ANOVA followed by Tukey’s HSD, naive vs. α-syn treated with quinpirole p = 0.9948, α-syn vs. α-syn treated with quinpirole p = 0.0003).

As a complementary approach, via a double-blinded experimental design, we used HPLC, as described in the Methods and in Figure 4, to measure extracellular dopamine level in the external milieu of neurons after quinpirole pretreatment (0.5 μM for 48 hours). HPLC analysis revealed a reduction in basal dopamine release in all quinpirole-treated experimental groups, with the largest fold decrease in α-syn overexpressing neurons (Figure 7 N, n = 3 from independent biological replicates, one-way ANOVA followed by Tukey’s HSD, α-syn vs. α-syn treated with quinpirole p = 0.0325 and naive vs. α-syn treated with quinpirole p = 0.9959;). Therefore, through pharmacological manipulation of D2 receptors, the α-syn dysregulation of dopamine transmission is potentially reversible (n = 3 each, from 3 independent biological replicates, one-way ANOVA followed by Tukey’s HSD, α-syn vs. α-syn treated with quinpirole p = 0.0325 and naive vs. α-syn treated with quinpirole p = 0.9959). The restoration of extracellular dopamine could be due to decreased neuronal activity, decreased dopamine synthesis, or both. Since we have already examined the former (Figure 7A-K), to test the possibility of decreased dopamine synthesis, we used HPLC to measure intracellular dopamine levels via a double-blinded experimental design (described in Methods). Intracellular dopamine levels in quinpirole-treated α-syn overexpressing neurons were significantly reduced compared to untreated α-syn overexpressing neurons (Figure 7O, n = 3 independent biological replicates, one-way ANOVA followed by Tukey’s HSD, α-syn vs. α-syn treated with quinpirole p = 0.0449 and naive vs. α-syn treated with quinpirole p = 0.6197) shown in Figure 4 (pink dotted line; n = 3 each, from 3 independent replicates, one-way ANOVA followed by Tukey’s HSD, α-syn vs. α-syn treated with quinpirole p = 0.0325 and WT vs. α-syn treated with quinpirole p = 0.9959). Since activation of D2R negatively regulates tyrosine hydroxylase^35,36,66,75,76,124,126,127^ and neuronal activity^33–36,38,39^, we then tested the hypothesis that reduced intra- and extracellular dopamine are, in part, due to decreased TH protein levels. Via a double-blinded experimental design, we utilized quantitative ELISA, as described in Figure 4, to measure TH levels. As shown in Figure 7P, TH protein level is decreased in quinpirole-treated, α-syn overexpressing neurons compared to untreated (0.5 μM for 48 hours; n = 3, one-way ANOVA followed by Tukey’s HSD, WT vs. α-syn treated with quinpirole p = 0.4288 and α-syn vs. α-syn treated with quinpirole p = 0.6809). The partial rescue of α-syn-induced neuronal dysregulation after D2R activation is consistent with neuroprotective properties of D2 receptors described previously^104,128–131^. It has been shown that D2 autoreceptors suppress dopamine synthesis through a negative feedback mechanism, and thus reduce oxidative stress caused by a high level of cytoplasmic dopamine^128–130^. In addition, consistent with our data, activation of D2 autoreceptors mediates neuroprotection by reducing neuronal excitability, cytoplasmic dopamine, and calcium levels^104,131^ that can restore the balance between energy income, expenditure, and its availability^112^. The data presented in this study provide a potential druggable target that may revert or prevent the untoward consequences of α-syn burden on dopamine neuronal viability and activity.

### Concluding remarks

In this study, we found α-syn overexpression dysregulates the structural and functional properties of dopaminergic neurons. The untoward consequences of increased α-syn likely cascade across the neuron, protracting the neuronal processes, increasing calcium burdens and biophysical properties of dopamine neurons as measured by increased burst firing activity. We found that the endogenous self-regulation of dopaminergic neurons fails to restrain the exacerbation of these phenotypes. Thereby, the signaling of these neurons in their networks becomes erratic, potentially creating avalanching neuronal dysfunction. The dysregulation of dopamine signaling within the brain therefore precedes neuronal demise. However, we show that these progressive dysregulations can be reversed through pharmacological manipulation.

D2 autoreceptors feedback mechanism is one of the main autoinhibitory mechanisms regulating dopamine neuronal activity^36,38,132^. We found D2 autoreceptors activity is diminished in α-syn overexpressing dopamine neurons, and prolonged incubation with a D2 receptor agonist, quinpirole (48 hours, 0.5 μM), nearly restored the firing activity to its canonical levels, reinstated intra and extracellular dopamine levels, and prevented losses in neuronal demise and structural neural complexity. Notably, D2 receptor agonists (full and partial) have attained FDA approval and have made their way into the clinic; however, these are often tested in late-stage PD. Our results suggest that the current treatment timeline may occur too late and that the efficacy of this strategy requires early intervention to reduce the rate of neuronal demise. Most crucially, our results suggest that neuronal loss is preventable and with future exploration across other mechanistic pathways in addition to D2 receptor pharmacology will reveal intersectional treatments that may have the capacity to ameliorate PD.

## Materials and Methods

### The experiments in this study are performed via a blinded or when possible doubleblinded experimental design

#### Reagents and chemicals

The source, catalogue number and concentration of reagents, antibodies and chemicals used in this study are outlined in Table 1. All viral vectors utilized in this study are listed in Table 2.

**Table 1:**

Primary Neuron Culture
| Chemical name | Concentration | Vendor | Catalog Number | Use/ Application |
| --- | --- | --- | --- | --- |
| NaCl | 116 mM | Sigma- Aldrich | S7653 | dissociation media |
| NaHCO<sub>3</sub> | 26 mM | Sigma- Aldrich | S5761 | dissociation media |
| NaH<sub>2</sub>PO<sub>4</sub> | 2 mM | Sigma- Aldrich | S9638 | dissociation media |
| D-glucose | 25 mM | Sigma- Aldrich | G8769 | dissociation media |
| MgSO<sub>4</sub> | 1 mM | Sigma- Aldrich | M7506 | dissociation media |
| Cysteine | 1.3 mM | Sigma- Aldrich | C7352 | dissociation media |
| Papain | 400 units/ml | Worthington Biochemical Corporation | LS003127 | dissociation media |
| Kynurenic acid | 0.5 mM | Sigma- Aldrich | K3375 | dissociation media |
| DMEM | 51.45% | Thermo Fisher Scientific | 11330032 | Glia media |
| Fetal Bovine Serum | 39.60% | Gemini | 100-106 | Glia media |
| Penicillin/Streptomycin | 0.97% | Thermo Fisher Scientific | 15-140-122 | Glia media |
| Glutamax 100X | 0.97% | Thermo Fisher Scientific | 35050061 | Glia media |
| Insulin (25mg/ml stock) | 0.08% | Sigma-Aldrich | I5500 | Glia media |
| Neurobasal-A | 96.90% | Thermo Fisher Scientific | 10888022 | DIV0 neuronal media |
| B27 Plus | 1.90% | Thermo Fisher Scientific | A3582801 | DIV0 neuronal media |
| GDNF | 0.15% | Sigma-Aldrich | SRP3200 | DIV0 neuronal media |
| Glutamax 100X | 0.97% | Thermo Fisher Scientific | 35050061 | DIV0 neuronal media |
| Kynurenic acid | 0.08% | Sigma-Aldrich | K3375 | DIV0 neuronal media |
| Neurobasal-A | 97.10% | Thermo Fisher Scientific | 10888022 | maintenance neuronal media |
| B27 Plus | 1.90% | Thermo Fisher Scientific | A3582801 | maintenance neuronal media |
| Glutamax 100X | 0.97% | Thermo Fisher Scientific | 35050061 | maintenance neuronal media |

Live Cell Imaging
| Chemical name | Concentration | Vendor | Catalog Number | Use/ Application |
| --- | --- | --- | --- | --- |
| NaCl | 126 mM | Sigma-Aldrich | S7653 | Calcium imaging ACSF |
| KCl | 2.5 mM | Sigma-Aldrich | P9541 | Calcium imaging ACSF |
| CaCl<sub>2</sub> | 2 mM | Sigma-Aldrich | 223506 | Calcium imaging ACSF |
| NaH<sub>2</sub>PO<sub>4</sub> | 1.25 mM | Sigma-Aldrich | 71505 | Calcium imaging ACSF |
| MgSO<sub>4</sub> | 2 mM | Sigma-Aldrich | M7506 | Calcium imaging ACSF |
| Dextrose | 10 mM | Sigma-Aldrich | D9434 | Calcium imaging ACSF |
| NaHCO<sub>3</sub> | 24 mM | Sigma-Aldrich | S5761 | Calcium imaging ACSF |
| NaCl | 92 mM | Sigma-Aldrich | S7653 | HEPES ACSF |
| KCl | 2.5 mM | Sigma-Aldrich | P9541 | HEPES ACSF |
| CaCl<sub>2</sub> | 0.5 mM | Sigma-Aldrich | 223506 | HEPES ACSF |
| NaH<sub>2</sub>PO<sub>4</sub> | 1.2 mM | Sigma-Aldrich | P5655 | HEPES ACSF |
| NaHCO<sub>3</sub> | 30 mM | Sigma-Aldrich | S5761 | HEPES ACSF |
| MgSO<sub>4</sub> | 10 mM | Sigma-Aldrich | M7506 | HEPES ACSF |
| D-glucose | 25 mM | Sigma-Aldrich | G8769 | HEPES ACSF |
| HEPES | 20 mM | Sigma-Aldrich | H3375 | HEPES ACSF |
| Na-L-ascorbate | 5 mM | Sigma-Aldrich | A4034 | HEPES ACSF |
| Na-pyruvate | 3 mM | Sigma-Aldrich | P5280 | HEPES ACSF |
| Thiourea | 2 mM | Sigma-Aldrich | T7875 | HEPES ACSF |
| Dopamine hydrochloride | 1 µM | Sigma-Aldrich | H8502 | Treatment |
| Quinpirole hydrochloride | 10 µM | Sigma-Aldrich | Q102 | Treatment |

Electrophysiology
| Chemical name | Concentration | Vendor | Catalog Number | Use/ Application |
| --- | --- | --- | --- | --- |
| NaCl | 126 mM | Sigma-Aldrich | S7653 | Electrophysiology ACSF |
| KCl | 2.5 mM | Sigma-Aldrich | P9541 | Electrophysiology ACSF |
| CaCl<sub>2</sub> | 2 mM | Sigma-Aldrich | 223506 | Electrophysiology ACSF |
| NaH<sub>2</sub>PO<sub>4</sub> | 1.25 mM | Sigma-Aldrich | 71505 | Electrophysiology ACSF |
| MgSO<sub>4</sub> | 2 mM | Sigma-Aldrich | M7506 | Electrophysiology ACSF |
| Dextrose | 10 mM | Sigma-Aldrich | D9434 | Electrophysiology ACSF |
| NaHCO<sub>3</sub> | 26 mM | Sigma-Aldrich | S5761 | Electrophysiology ACSF |
| Potassium gluconate | 120 mM | Sigma-Aldrich | P1847 | Electrophysiology ACSF |
| MgCl<sub>2</sub> | 2 mM | Sigma-Aldrich | M8266 | Electrophysiology ACSF |
| HEPES | 10 mM | Sigma-Aldrich | H3375 | Electrophysiology ACSF |
| EGTA | 0.1 mM | Sigma-Aldrich | E3889 | Electrophysiology ACSF |
| ATPNa<sub>2</sub> | 2 mM | Sigma-Aldrich | A2383 | Electrophysiology ACSF |
| GTPNa | 0.25 mM | Sigma-Aldrich | A2383 | Electrophysiology ACSF |
| Dopamine hydrochloride | 1 µM | Sigma-Aldrich | H8502 | Electrophysiology ACSF |
| Quinpirole hydrochloride | 10 µM | Sigma-Aldrich | Q102 | Electrophysiology ACSF |

Biochemical Assays
| Chemicals | Concentration | Vendor | Catalog Number | Use/ Application |
| --- | --- | --- | --- | --- |
| Phosphate Buffered Saline (PBS) | 1X | Prepared as needed on-site | N/A | ICC |
| Triton X-100 | 0.005 | Thermo Fisher Scientific | bp151-100 | ICC |
| Normal Goat Serum (NGS) | 0.1 | Lampire Biological Products | 7332500 | ICC |
| Paraformaldehyde | 0.04 | Electron Microscopy Sciences | 157-4-100 | ICC |
| Na<sub>2</sub>CO<sub>3</sub> | 28.3 mM | Sigma-Aldrich | D6546 | ELISA Coating Buffer |
| NaHCO<sub>3</sub> | 71.42 mM | Sigma-Aldrich | S5761 | ELISA Coating Buffer |
| Glycerol | 10% (v/v) | Sigma-Aldrich | G5516 | BufferD Lysis Buffer |
| NaCl | 125 mM | Sigma-Aldrich | S7653 | BufferD Lysis Buffer |
| EDTA | 1 mM | Sigma-Aldrich | E9884 | BufferD Lysis Buffer |
| EGTA | 1 mM | Sigma-Aldrich | 3777 | BufferD Lysis Buffer |
| NaCl | 150 mM | Sigma-Aldrich | S7653 | Biotinylation Buffer |
| CaCl<sub>2</sub> | 2 mM | Sigma-Aldrich | 449709 | Biotinylation Buffer |
| Triethanolamine | 10 mM | Sigma-Aldrich | 90279 | Biotinylation Buffer |
| Sulfo-NHS-SS Biotin | 1.5 mg/mL | ThermoFisher Pierce | 22331 | Biotinylation Buffer |
| Monomeric Avidin Resin | n/a | ThermoFisher Pierce | 53146 | Biotinylation Buffer |
| Fat-free milk | 1% or 5% | Carnation | N/A | WB/ELISA |
| TMB Substrate | Stock | ThermoFisher | 34028 | ELISA |
| H<sub>2</sub>SO<sub>4</sub> | 2N | Sigma-Aldrich | 339741 | ELISA |
| Tween-20 | 0.002 | ThermoFisher | MP1Tween201 | WB/ELISA: TBS-T |
| Protease Inhibitor | 1x | Millipore | 539191 | WB/ELISA: Cell lysis |
| DC Protein assay | N/A | Biorad | 5000112 | Protein assay |
| Immulon 4HBX | N/A | ThermoFisher | 3855 | ELISA |
| NaCl | 118 mM | Sigma-Aldrich | S7653 | KH Buffer (HPLC) |
| KCl | 4.7 mM | Sigma-Aldrich | P9541 | KH Buffer (HPLC) |
| CaCl<sub>2</sub> | 1.25 mM | Sigma-Aldrich | 223506 | KH Buffer (HPLC) |
| KH<sub>2</sub>PO<sub>4</sub> | 1.2 mM | Sigma-Aldrich | P5655 | KH Buffer (HPLC) |
| MgSO<sub>4</sub> | 1.2 mM | Sigma-Aldrich | M7506 | KH Buffer (HPLC) |
| D-glucose | 11 mM | Sigma-Aldrich | G8769 | KH Buffer (HPLC) |

| L-Ascorbic acid | 0.5 mg/ml | Sigma-Aldrich | A5960 | KH Buffer (HPLC) |  |
| --- | --- | --- | --- | --- | --- |
| NaHCO<sub>3</sub> | 25 mM | Sigma-Aldrich | S5761 | KH Buffer (HPLC) |  |
| Antibodies |  |  |  |  |  |
| Primary Antibody | Host Species / Isotype | Concentration | Vendor | Catalog Number | Use |
| Tyrosine Hydroxylase | Rabbit / Polyclonal | 1:500 | EMD Millipore | AB152 | ICC/WB |
| α-synuclein (syn211) | Mouse/ IgG1 | 1:500 | Abcam | AB80627 | ICC |
| Dopamine Transporter (DAT) | Rat | 1:500 | Millipore Sigma | AB5802 |  |
| Calbindin | Rabbit / Polyclonal | 1:500 | EMD Millipore | ABN2192 |  |
| Green Fluorescent Protein (GFP) | Rabbit / IgG | 1:500 | ThermoFischer Scientific | A-11122 | ICC |
| Human α-syn (C-terminal regions, 130-140) (94-3A10) | Mouse/ monoclonal IgG1 | 1:1000 | Gifted from Dr. Giasson | N/A | WB |
| Actin C4 | Mouse / Monoclonal | 1:1000 | EMD Millipore |  | WB |
| D2 | Mouse | 1:1000 | Neuromab | 75-230 | WB/ICC |
| D2 | Rabbit |  | Millipore |  | WB |
| TH | Monoclonal/Mouse | 1:1000 | EnCor | MCA-4H2 | ELISA |
| TH (Biotin conjugate) | Polyclonal/Rabbit | 1:6000 | EnCor | RPCA-TH | ELISA |
| Biotin (HRP conjugate) | N/A (Avidin) | 1:200 | Vector Labs | A2004 | ELISA |
| Conjugated (or not) Secondary Antibody | Host species / Target species | Concentration | Vendor | Catalog Number | Use |
| Alexa Fluor 568 | Goat / mouse | 1:500 | Life Technologies | A-21124 | ICC |
| Alexa Fluor 647 | Goat / mouse | 1:500 | Life Technologies | A-21242 | ICC |
| Alexa Fluor 488 | Goat / Rabbit | 1:500 | Life Technologies | A-11006 | ICC |
| HRP | Goat / Mouse | 1:2000 | Jackson Immuno Research Labs | AB-10015289 | WB |
| HRP | Goat / Rabbit | 1:2000 | Jackson Immuno Research Labs | AB_2307391 | WB |
| IRDye® 800CW | Goat / Mouse | 1:15000 | Li-cor | 926-32210 | WB |

**Table 2:**

| Table 2: |  |  |  |  |  |
| --- | --- | --- | --- | --- | --- |
| Virus |  |  |  |  |  |
| Viral Vectors (AAV) | Concentration | Vendor | Catalog Number | Capsid Serotype | Other |
| TH-α-syn | $3.26 \times 10^7$ units/μl | Dr. Giasson lab | N/A | AAV 1 | |
| TH-GFP | $3.26 \times 10^7$ units/μl | Dr. Giasson lab | N/A | AAV 1 | |
| LoxP - tdTomato | $3.26 \times 10^7$ units/μl | Addgene | 28306 | AAV 1 | |
| Virus |  |  |  |  |  |
| Viral Vectors (AAV) | Concentration | Vendor | Catalog Number | Capsid Serotype | Other |
| TH-α-syn | 3.26×10<sup>7</sup> units/μl | Dr. Giasson lab | N/A | AAV 1 |  |
| TH-GFP | 3.26×10<sup>7</sup> units/μl | Dr. Giasson lab | N/A | AAV 1 |  |
| LoxP - tdTomato | 3.26×10<sup>7</sup> units/μl | Addgene | 28306 | AAV 1 |  |

#### Animals

All experiments were approved by the Institutional Animal Care and Use Committee at University of Florida. Mice were housed in the animal care facility at the University of Florida, 2-4 per cage with food and water available ad libitum in the home cage. The room was maintained under 12-hour light/dark cycle. Wild-type C57BL/6J mice, or DAT^IRES*cre*^ and Ai95(RCL-GCaMP6f)-D (Ai95D) knock-in mice were obtained from The Jackson Laboratory (Stock number: 006660 (DAT^IRES*cre*^), 024105 (Ai95D), Bar Harbor, ME, USA). WT pups, or pups expressing GCaMP6f in dopamine neurons were used for this study. Mice of both sexes were used.

#### AAV1-TH-α-synuclein and AAV1-TH-GFP generation

The pAAV2.5-THP-GFP plasmid was purchased from Addgene (#80336)^133^. Human α-synuclein cDNA from a pcDNA3.1 plasmid was restriction digested with EcoRI and HindIII (NEB), purified and ligated in the same sites of the pAAV2.5-THP backbone to generate the pAAV2.5-TH-α-syn plasmid. Purified pAAV2.5TH-α-syn and pAAV2.5-TH-GFP vectors were utilized to prepare active AAV (capsid 1) using a HEK293T-based transfection method followed with iodixanol gradient purification as previously described^134^ and these virus were termed AAV1-TH-α-syn and AAV1-TH-GFP. Genomic titer of each virus was assayed by quantitative PCR as previously described^134^.

#### Primary Neuronal Culture

Primary culture was prepared as previously described, with small distinctions^50^. Briefly, acutely dissociated mouse midbrains from 0-2-day old male and female pups were isolated and incubated in dissociation medium at 35-37 °C under continuous oxygenation for 60-90 min. Dissociated cells were triturated with pipettes of decreasing bore size (including a punctured fire-polished pipette), then pelleted by centrifugation at 1500 rpm for 3-5 min, and resuspended and plated in glial medium (Table 1). Cells were plated at a density of 100,000 cells/coverslip, on 12 mm coverslips coated with 0.1 mg/ml poly-D-lysine and 5 μg/ml laminin and maintained in neuronal media. After 2 hours, cells were supplemented with neuronal media (DIV0 composition). Every 4 days, half of the media was replaced with fresh media. On DIV5, cultures were transduced with the desired AAV1 (see Table 2). The experiments described in this study were performed on DIV9-11. Reagents and chemicals utilized for midbrain neuronal culture are listed in Table 1.

#### Electrophysiology

Spontaneous firing activity of midbrain dopamine neurons was examined via whole cell current clamp recordings as previously described^46,48,51^. The neurons were continuously perfused with ACSF (composition described in Table 1) equilibrated with 95% O2/5% CO2; pH was adjusted to 7.4 at 37 °C. Patch electrodes were fabricated from borosilicate glass (Cat. No. 1B150F-4,1.5 mm outer diameter; World Precision Instruments, Sarasota, FL) with the P-2000 puller (Sutter Instruments, Novato, CA). The tip resistance was in the range of 3-5 MΩ. The electrodes were filled with a pipette solution containing (in mM): 120 potassium gluconate, 20 KCl, 2 MgCl2, 10 HEPES, 0.1 EGTA, 2 ATP, and 0.25 GTP, with pH adjusted to 7.25 with KOH. All experiments were performed at 37 °C. To standardize action potential (AP) recordings, neurons were held at their resting membrane potential (see below) by DC application through the recording electrode. Action potential was recorded if the following criteria were met: a resting membrane potential more polarized than −35 mV and an action potential peak amplitude greater than 60 mV. Action potential half-width was measured as the spike width at the half-maximal voltage using Clampfit 10 software (Molecular Devices LLC, San Jose, CA). Steady-state basal activity was recorded for 2-3 min before bath application of the drug. Each coverslip was used for only one recording, this is specifically important for experiments involving drug application. The spontaneous spike activity of midbrain dopamine neurons was obtained by averaging 1 min interval activities at baseline and after 3-5 min of drugs.

#### Live Cell Calcium Imaging

*These experiments were performed via a blinded experimental design*. Live cell calcium imaging and analysis are described previously^50^. Briefly, naive (non-transduced) and α-syn overexpressing (transduced with AAV1-TH-α-synuclein) midbrain neuronal cultures were imaged with a Nikon Eclipse FN1 upright microscope (Nikon Instruments, Melville, NY). A Spectra X (Lumencor, Inc, Beaverton, OR) was used to stimulate GCaMP6f (λex = 470 nm) fluorescence through a custom quad-pass filter (Chroma Technologies, Battleboro, VT), and emission was filtered through visible spectra bandpass filter. Experiments were performed under gravity perfusion of artificial cerebral spinal fluid (ACSF, Table 1). The average fluorescence of the first 60s recording is defined as baseline. After baseline imaging, vehicle (ACSF), 1 μM DA, or 10 μM quinpirole was administered via a perfusion system (flow rate of 2 ml/min) and recorded for additional 2 minutes. Neuronal somatic regions of interest (ROI) were auto detected for the cell body of each individual neuron. Background fluorescence was subtracted from each frame. Fold fluorescence change from baseline was calculated and plotted against time. Each coverslip was used for only one recording, this is specifically important for experiments involving drug application. All resources and reagents used for live cell calcium imaging experiments are listed in Table 1.

#### Live cell confocal imaging using GRAB_DA4.4-_ expressing HEK 293 cells to measure *extracellular dopamine*

*These experiments were performed via a double-blinded experimental design*. GRAB_DA4.4_ is a genetically encoded fluorescent dopamine sensor, engineered by coupling a conformationally-sensitive circular-permutated EGFP (cpEGFP) to D2 receptor. In GRAB_DA4.4-_expressing HEK293 cells, dopamine binding to the sensor induces a conformational change which results in a robust increase in fluorescence signal via a concentration-dependent manner^135^. Flp-In™ 293 T-Rex stable cell lines exhibiting tetracycline-inducible expression of the GRAB_DA4.4_ dopamine sensor was generously gifted by Dr. Ulrik Gether. The GRAB_DA4.4-_ expressing HEK 293 cells were maintained in DMEM supplemented with 10% FBS and 100 unitsIml PenicillinIStreptomycin. Selection pressure for GRAB_DA4.4_ expressing cells was maintained with media containing Hygro-B (1 mgIml) and Blasticidin (0.015 mgIml). The cells were plated on coverslips in media containing tetracycline (1:1000) to induce expression for 12-24 hours before live cell imaging under three conditions: 1) only GRAB_DA4.4-_expressing HEK293 cells, to measure constitutive fluorescent signal, 2) GRAB_DA4.4_ cells added to tdTomato expressing dopaminergic neurons, 3) GRAB_DA4.4_ cells added to tdTomato expressing dopamine neurons overexpressing α-syn. The tdTomato transduction (λex = 560 nm) is used for a better identification of soma and dendritic field and no fluorophore overlap (i.e. excitation and emission of tdTomato does not bleed through GFP channel). Imaging was performed using a Nikon Eclipse FN1 upright microscope (Nikon Instruments, Melville, NY). A Spectra X (Lumencor, Inc, Beaverton, OR) was used to stimulate GFP (λ_ex_ = 470 nm), tdTomato (λ_ex_ = 560 nm) fluorescence through a custom quad-pass filter (Chroma Technologies, Battleboro, Vermont). Emission was filtered through visible spectra bandpass filter. Regions of interest (ROI) were autodetected via NIS Elements software (Nikon Instruments, Melville, NY). cpEGFP signal was background subtracted using an ROI in an adjacent area of the image devoid of cells or debris.

#### Generation of standard curve

To generate a standard curve, the baseline fluorescence signal (F_c_) which is the constitutive fluorescent signal in the absence of extracellular dopamine was recorded. Changes in fluorescence signal after adding various dopamine concentrations (1-20 nM) were plotted against dopamine concentration.

#### Measurement of basal dopamine release

GRAB_DA4.4_ cells are co-cultured with tdTomato expressing dopamine neurons (DAT^IRES*cre*^-LoxP-tdTomato) containing endogenous α-syn or its overexpression 20-24 hours prior to live cell confocal imaging. At the beginning of each experiment, the constitutive GRAB_DA4.4_ fluorescence signal (F_c_) of the cells that are plated in similar conditions sans neurons was obtained. To compare baseline dopamine release amongst the experimental groups, the average ratio of fluorescence signal of cells adjacent to the soma and neuronal processes to the average ratio of fluorescence signal of GRAB_DA4.4_ cells (only) were calculated (F_baseline_ = (F_GRAB DA4.4 cells grown with neurons_ - F_c_) I F_c_).

#### Visualization and quantification of real-time dopamine release following KCl stimulation

20-24 hours prior to live cell confocal imaging, GRAB_DA4.4_ cells are cocultured with tdTomato expressing dopamine neurons containing endogenous α-synuclein or its overexpression. GRAB_DA4.4_ fluorescent signal around the soma and neuronal processes were measure before (F_baseline_) and following KCl-stimulation (90 mM) of dopamine release^57^. Average fluorescence signal of cells adjacent to the soma and neuronal processes before and after KCl were calculated (ΔF/F = (F_stimulated_ - F_baseline_) *I* F_baseline_).

#### Immunocytochemistry (ICC)

*These experiments were performed via a double-blinded experimental design.* On DIV9-11, naive (non-transduced) and α-syn overexpressing neuronal cultures were fixed with 4% paraformaldehyde (PFA) in PBS for 30 min at room temperature (RT), followed by blocking, permeabilization and overnight incubation (at 4 °C) with primary antibodies diluted in blocking buffer, followed by three 20-min phosphate buffer solution (PBS) washes. Then, a 1-hour incubation in blocking buffer with Alexa- Fluor conjugated secondary antibodies at RT, followed by three 20-min washes and an overnight PBS wash at RT. Coverslips were mounted on slides using Fluoromount-G. Images were captured on a Nikon A1 laser-scanning confocal microscope (20X or 40X oil-immersion objective). Reagents and chemicals utilized for ICC are listed in Table 1.

#### Western Blot Analysis

*These experiments were performed via a double-blinded experimental design*. For detection of endogenous and human α-syn total cell lysate of neurons transduced with AAV1-TH-α-syn (n = 4) or naive (non-transduced) neurons (n = 4) were used, as described previously^136^ Briefly, the cells were harvested in 200 μL of 2% SDS buffer, protein concentrations were determined using the bicinchoninic acid assay (Pierce), and further diluted in sample buffer (10 mM Tris, pH 6.8, 1 mM EDTA, 40 mM DTT, 0.005% Bromophenol Blue, 0.0025% Pyronin Yellow, 1% SDS, 10% sucrose). Following harvest of total cell lysate, samples were heated to 100 °C for 10 minutes prior to SDS-PAGE (13% polyacrylamide gels, 10 μg lysate per well) followed by electrophoretic transfer onto 0.2 μm nitrocellulose membranes as previously described^136^ Membranes were incubated with block solution (5% milk in TBS) for 1 h, then with primary antibodies (in block solution) overnight at 4°C. Membranes were washed with TBS-T and incubated with goat anti-mouse or anti-rabbit secondary antibodies conjugated to horseradish peroxidase (Jackson Immuno Research Labs, Westgrove, PA) diluted in block solution at room temperature for 1 hour. Immunoreactivity was assessed using Western Lightning-Plus ECL reagents (PerkinElmer, Waltham, MA) followed by chemiluminescence imaging (Genegnome XRQ, Syngene, Frederick, MD). For α-syn detection we used 94-3A10 antibody, which is a mouse monoclonal antibody raised against C-terminal regions (130-140) of α-syn^137^ For loading control, we used mouse monoclonal anti-actin C4 antibody (EMD Millipore). For tyrosine hydroxylase detection, an affinity purified rabbit antibody AB152 (EMD Millipore) was used.

#### Biotinylation Assay

*These experiments were performed via a double-blinded experimental design*. α-Syn overexpressing neuronal cultures and non-transduced neuronal cultures were washed three times with cold PBS and incubated with sulfo-NHS-biotin (1.5 mg/ml; ThermoFisher Pierce, 21331) for 30 minutes at 4 °C with rocking. Remaining sulfo-NHS-biotin was quenched with cold Quenching Solution (Glycine 50 mM in PBS), followed by three washes with cold PBS^138,139^. Cells were lysed in BufferD lysis buffer (10% Glycerol, 125 mM NaCl, 1 mM EDTA, 1 mM EGTA, pH 7.6) containing 1% Triton-X100 and protease inhibitor cocktail (Millipore, 539131) for 1 hour at 4 °C with rocking, followed by centrifugation for 15 minutes at 12,000 g. The supernatants were divided into three portions - 25 μl for protein quantification, 200 μl for incubation with Avidin, with the remainder for whole lysate. After equilibrating monomeric UltraLink Avidin (ThermoFisher Pierce, 53146) twice with 1 ml BufferD, 40 μl of 50% bead slurry were added to 200 μl lysate and incubated at 4 °C for 1 hour with rotation. Supernatant was retained as cytoplasmic fraction, and beads were washed three times with 1 ml BufferD, eluted with 40 μl Laemmli Sample Buffer 4x (containing 10% beta-mercaptoethanol) at 37 °C for 30 minutes and separated by 10% SDS-PAGE, transferred to 0.45 μm nitrocellulose and probed with antibodies against proteins of interest (see Table 1). Fluorescent images were analyzed using ImageJ (NIH) to measure band optical density. Values were normalized to total protein per lane. Beta-tubulin (Aves, TUJ) was probed to demonstrate membrane fraction isolation during biotinylation.

#### ELISA Quantification of Tyrosine Hydroxylase (TH)

These experiments were performed via a double-blinded experimental design.

#### Cell lysis

For total protein quantification via ELISA, neuronal cultures were washed three times with cold PBS, then lysed in BufferD lysis buffer (10% Glycerol, 125 mM NaCl, 1 mM EDTA, 1 mM EGTA, pH 7.6) containing 1% Triton-X100 and protease inhibitor cocktail (Millipore, 539131) for 1 hour at 4 °C with rocking, followed by centrifugation for 15 minutes at 12,000g. Samples were denatured in Laemmli Sample Buffer 4x (containing 10% beta-mercaptoethanol) at 37 °C for 30 minutes and separated by 10% SDS-PAGE, transferred to 0.45 μm Nitrocellulose and probed with antibodies against proteins of interest. Values were normalized to total protein per lane.

#### TH ELISA

Antibodies and concentrations used are given in Table 1. In brief, Immulon 4 HBX High-Binding 96 well plates were coated with 100 μl per well of 1:1,000 dilution of mouse anti-TH (EnCor, MCA-4H2) in coating buffer (28.3 mM Na_2_CO_3_, 71.42 mM NaHCO_3_, pH 9.6) for 20 hours at 4 °C. Edge lanes 1 and 12 were left empty. Wells were blocked with 5% fat free milk in 1x TBS (pH 7.4) for 1 hour at room temperature on an orbital shaker set to 90rpm.

#### Generation of standard curve

To produce a standard curve, two standard curve lanes were generated, with six serial dilutions, beginning at 10 ngIml and 1 ngIml in TBS-T containing 1% fat free milk (with the last well in each standard curve lane left with incubation buffer only as a blank. Remaining wells were incubated in duplicate with lysates from cells of interest. Incubation was completed for 20 hours at 4 °C on an ELISA shaker set to 475 rpm. After each well was washed and aspirated 6 times with TBS-T, anti-TH rabbit (EnCor, RPCA-TH) conjugated to biotin was dilute 1:6,000 from a stock concentration of 1.65 mgIml in TBS-T with 1% fat-free milk and incubated for 1 hour at room temperature at 425 rpm. 100 μl Avidin-HRP (Vector labs, A-2004), diluted 1:2,500 in TBS-T with 1% fat-free milk, was added to each well following wash, and incubated for 1 hour at room temperature at 425 rpm. Following final washes, 150 μl room temperature TMB-ELISA reagent (Thermo Fisher, 34028) was added to each well. The reaction was allowed to continue for 20 minutes, protected from light, and stopped by addition of 50 μl 2N H2SO4. The plate was immediately read at 450 nm. Duplicate standard and sample wells were averaged, and background-subtracted based on blank wells. TH Concentration was calculated using a quadratic curve equation calculated in Graphpad Prism 8, then normalized to total protein concentration per sample as calculated using Lowry Assay. Final TH values shown are presented as pg THImg total protein after multiplication of the nanogram TH value by 1,000 to show TH as picogram THImilligram total protein.

#### HPLC

*These experiments were performed via a double-blinded experimental design.* Midbrain primary cultures were incubated in KH buffer (Table 1), at 37 °C, for 1 hour before collecting intracellular and extracellular milieu for HPLC analysis^140,141^. For extracellular milieu, KH buffer incubated with neurons was collected, treated with 1M perchloric acid and snap frozen for analysis. For intracellular milieu, coverslips were washed with KH buffer, scraped, and treated with 1M perchloric acid, before sonicating. Then sample was centrifuged at 12000 rpm at 4°C for 10 min, and supernatant snap-frozen in liquid nitrogen for analysis. The pellet was resuspended using 0.2 NaOH and RIPA buffer^142^ for protein quantification via Lowry assay. Samples were centrifuged at 16,000 g for 15 min (4°C) and the supernatant was filtered through a 0.2 mm pore membrane (Nanosep with 0.2 mm bioinsert, Pall Life Sciences) and 15 μl of the supernatant was injected directly into an HPLC-ECD (HTEC-510; Eicom). Dopamine was separated on a CAX column (EICOMPAK 2.0 id x 200 mm) maintained at 35°C. The mobile phase consisted of 70% 0.1 M ammonium acetate buffer (pH 6.0) containing sodium sulfate (0.025 M), EDTA-2Na (50 mg/l) and 30% methanol at a flow rate of 250 μl/min. An electrochemical detector that used a glassy working electrode (+ 450 mV) against a silver-silver chloride reference electrode (WE-3G; Eicom) was used to quantify dopamine in the samples. A dopamine standard was used to identify and quantify the dopamine concentration in the samples.

#### Morphometric Analysis

*These experiments were performed via a double-blinded experimental design*. Control (DAT^IRES*cre*^) neurons and α-syn overexpressing neurons (DAT^IRES*cre*^/α-syn) were transduced with AAV1-LoxP-tdTomato (Addgene) on DIV5. Neurons were fixed with 4% paraformaldehyde for 30 minutes at room temperature on DIV10, and coverslips were mounted using Fluoromount-G and allowed to dry. Alternatively, neurons were fixed, co-immunolabeled with TH and GFP antibodies, and mounted for imaging, as described above. Images were captured on a Nikon A1 laserscanning confocal microscope (visualized through a 20X oil-immersion objective). Images of neurons with minimal interference from neighboring neurons were analyzed in ImageJ (FIJI) and converted to 8-bit binary images after threshold adjustment. Sholl analysis plugin was used to draw concentric circles starting from 15 μm followed by 5 μm successive shells in order to identify the number of intersections along the radii^94,143,144^. Sholl analysis was performed and the number of intersections were plotted (Figure 6). Cell area measurements were attained by manually drawing ROI around cell soma using the free polygon selection tool in ImageJ. ROIs were drawn to encompass the complete projection area of the cell and selection was finalized by convex hull to attain final projection area measurement. Results obtained were plotted to analyze complexity of morphology in non-transduced controls and *α-synuclein-expressing* neurons.

#### Statistical Analysis

Data analysis was performed using Graphpad Prism version 8.02 and MATLAB version 2020a. Student’s t-test, linear regression, one-way, two-way, or repeated measures ANOVA were used where appropriate and corrected for multiple comparisons. Significance of *P*<0.05 was considered statistically significant. Data are presented with mean and standard error unless otherwise stated.

## Contributions

H.K. and B.G. conceived and supervised the study. H.K., B.G., A.D., D.R.M, A.G., Z.A.S. and J.J.L. designed and planned the experiments. A.D., D.R.M., F.S., M.L., A.G., S.H., Z.A.S., and J.J.L. collected the data. A.D., D.R.M., S.V., A.R.A., J.A., and C.A.H. analyzed the data. H.K., B.G., A.D., and D.R.M. interpreted the data with comments from N.U. H.K., B.G., A.D., and D.R.M. prepared the manuscript. B.U. and A.M.B. revised the work.

---

**Supplemental Figure 1:**
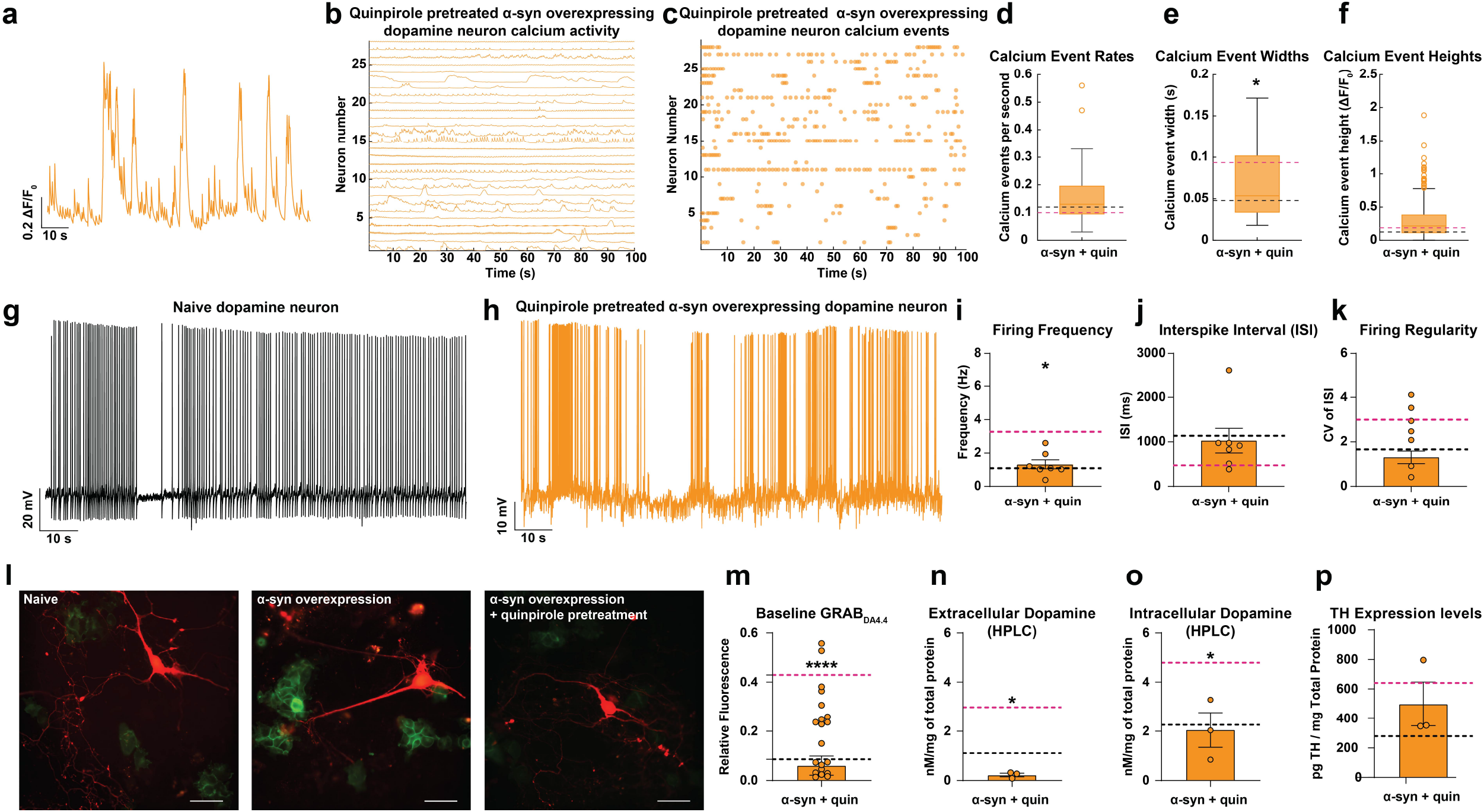
AAV transduction of TH-dependent α-synuclein produces robust overexpression. Quantification of multiple blots illustrating increased α-synuclein levels in naive and transduced cultures (n = 4, two-tailed t-test, p = 0.005).

**Supplemental Figure 2:**
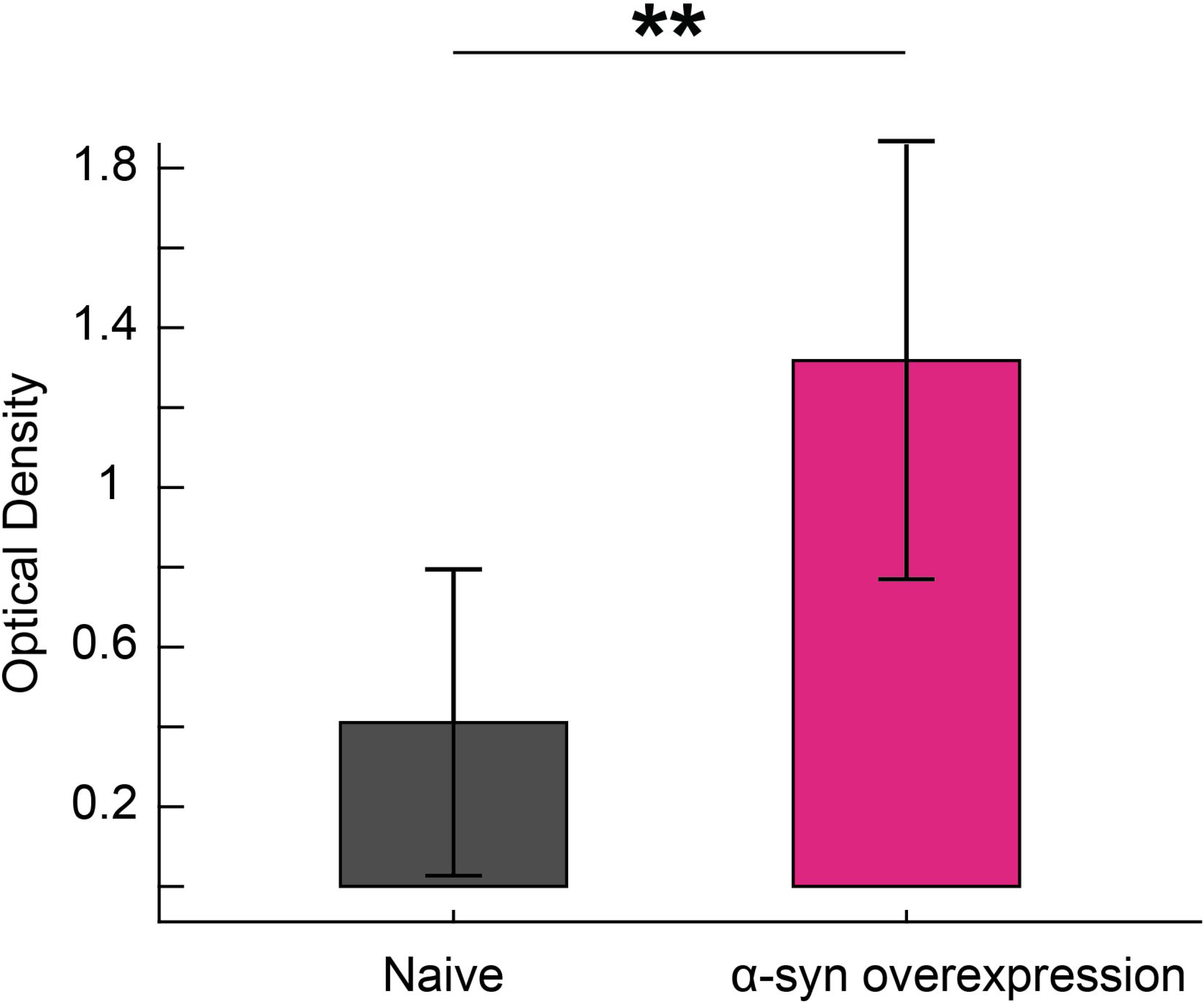
Overexpression of control vector (AAV-TH-GFP) does not affect neural response. (A,B) Neurons overexpressing GFP do not produce responses different from naive neurons in response to either dopamine (A) or quinpirole (B) (Dopamine: n = 11-17, one-way ANOVA followed by Tukey’s HSD, naive vs. α-synuclein overexpression, p = 0.0035, naive vs. GFP p = 0.2927 and α-syn vs. GFP p = 0.0001, from five independent replicates, Quinpirole: n = 11-13, one-way ANOVA followed by Tukey’s HSD, naive vs. α-syn overexpression, p = 0.0318, naive vs. GFP p = 0.8178, and α-syn vs. GFP p = 0.0057, from five independent replicates).

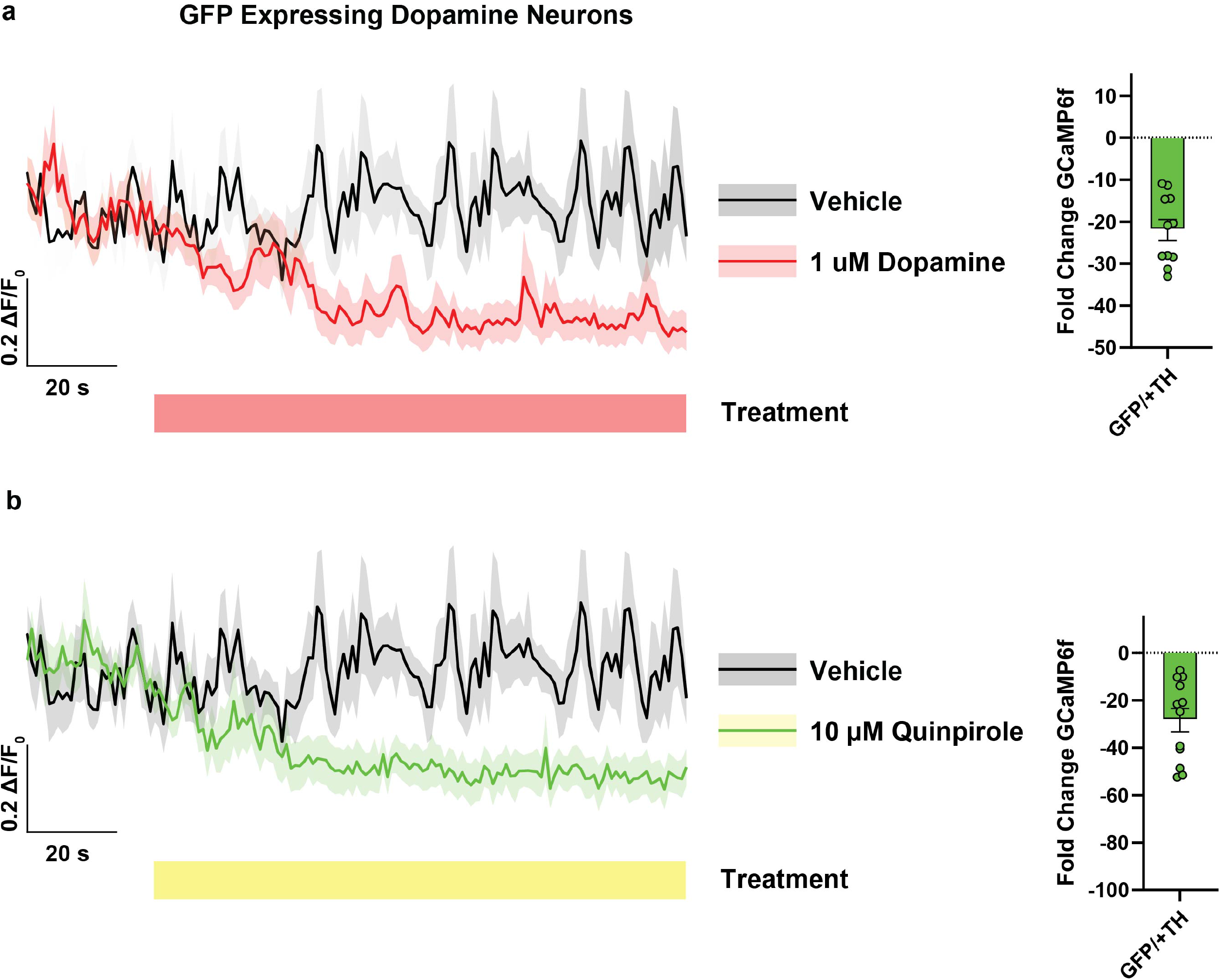

